# ENPP3 expressed by HER2-positive breast cancer cells is associated with good prognosis by restraining epithelial-to-mesenchymal phenotype

**DOI:** 10.64898/2026.08.31.747828

**Authors:** Roberta Bongiorno, Maria Teresa Majorini, Valeria Figà, Davide Pernici, Matteo Dugo, Antonino Belfiore, Paolo Baili, Valeria Cancila, Elena Jachetti, Giancarlo Pruneri, Serenella Pupa, Loris De Cecco, Tiziana Triulzi, Claudio Tripodo, Mario Paolo Colombo, Claudia Chiodoni, Daniele Lecis

**Affiliations:** Molecular Immunology Unit, Department of Experimental Oncology, Fondazione IRCCS Istituto Nazionale dei Tumori di Milano, Milan, Italy; Breast Cancer Unit Clinical Translational and Immunotherapy Research, Department of Medical Oncology, IRCCS Ospedale San Raffaele, Milan, Italy; Department of Diagnostic Pathology and Laboratory Medicine, Fondazione IRCCS Istituto Nazionale Dei Tumori di Milano, Milan, Italy; Department of Epidemiology and Data Science, Data Science Unit, Fondazione IRCCS Istituto Nazionale dei Tumori di Milano, Milan, Italy; Tumor Immunology Unit, Department of Health Sciences, University of Palermo, Palermo, Italy; Microenvironment and Biomarkers of Solid Tumors Unit, Department of Experimental Oncology, Fondazione IRCCS Istituto Nazionale Dei Tumori di Milano, Milan, Italy; Integrated Biology of Rare Tumors, Department of Experimental Oncology, Fondazione IRCCS Istituto Nazionale dei Tumori di Milano, Milan, Italy; IFOM ETS – The AIRC Institute of Molecular Oncology, Milan, Italy

**Keywords:** prognostic marker, ectonucleotide pyrophosphatase/phosphodiesterase

## Abstract

**Background:** Ectonucleotide pyrophosphatase/phosphodiesterase 3 (ENPP3/CD203c) is largely studied as a marker of mast cells and basophils. By depleting extracellular ATP, it prevents excessive activation of mast cells and basophils, hence reducing inflammation and allergic reactions. Recent findings have also shown that Enpp3 can deplete cGAMP, another molecule involved in STING activation and IFN-mediated pro-inflammation. Little is still known regarding the role of Enpp3 in non-immune cells although a few reports have described its expression in healthy tissues and tumors.

**Methods:** *In silico* analysis were performed to investigate the expression levels and the prognostic value of Enpp3 in breast cancer, together with ovarian, prostate and colon carcinoma. ENPP3 expression was evaluated in formalin-fixed, paraffin-embedded tumor samples of breast cancer patients by immunohistochemistry, and in mouse mammary cancer cell lines by western blots. Cells were treated with EGFR ligands to stimulate the EGFR/HER2 axis. A mouse-derived mammary cancer cell line was engineered by CRISPR/Cas9 to introduce a GFP sequence under the control of the *Enpp3* promoter. GFP-positive and - negative cells were sorted and analyzed by gene expression profiling to identify genes and pathways associated with *Enpp3* expression. Finally, wild type and Enpp3 knockout cells were injected in the fat pad of Wsh mice, which do not have mast cells, to evaluate the growth of the tumors which were further analyzed by immunohistochemistry.

**Results:** We provide evidence that HER2-positive cells express higher levels of ENPP3 in samples of breast cancer patients. Moreover, in *vitro* models confirmed that HER2 expression and EGFR stimulation result in up-regulation of Enpp3. We identified pathways that can concur to *Enpp3* expression and showed that i*n vivo* the absence of Enpp3 promotes tumor growth and development of tumors with a marked epithelial-to-mesenchymal phenotype. Finally, in a small cohort of HER2-positive breast cancer patients, we found that ENPP3 expression correlates with increased relapse-free survival.

**Conclusions:** Despite its potential immunosuppressive role, our findings support the notion that ENPP3 expression is promoted by HER2 in breast cancer, and that it is endowed with a positive prognostic value.

## Background

Ectonucleotide pyrophosphatase/phosphodiesterase 3 (ENPP3), also known as CD203c, is a type II transmembrane glycoprotein which belongs to the family of ENPP ectoenzymes. ENPPs hydrolyze extracellular ATP and ADP, together with other nucleotide substrates. In this way, they regulate purinergic signals and affect many processes, ultimately regulating inflammation, tissue remodeling, and especially immune cell activation^1^. In particular, ENPP3 is endowed with phosphodiesterase and pyrophosphatase activity, and it is able to hydrolyze ATP to produce AMP and pyrophosphate, which are involved in immunosuppression^2,3^ and inhibition of mineralization^4^, respectively. Hence, ENPP3, by degrading nucleotides, can affect many physiological and pathological cell features.

The vast majority of works describing ENPP3 activity have focused on its immunological role. It is considered a marker of mast cells and basophils^5^, which rapidly upregulate ENPP3 upon IgE receptor engagement^6,7^. ENPP3 expression is exploited as a marker for mast cell activation and degranulation in allergic diseases^8,9^. In this context, ENPP3 attenuates inflammation by reducing mast cell activation^10^, and preventing the release of pro-inflammatory mediators such as histamine and cytokines^8^. Hence, ENPP3 can contribute to immune response and shaping of the immune contexture.

Although the role of ENPP3 in mast cells and basophils has been extensively studied, both for its activity and as a marker in allergic reaction, little is still known regarding ENPP3 function in non-immune cells and especially in tumors. ENPP1, which shares a high homology with ENPP3, has been shown to play a crucial role in tumor progression and metastasis^11^. This activity stems from its capability to hydrolase cGAMP, therefore negatively regulating STING-mediated interferon signaling and immune activation^11,12^. Consequently, ENPP1 is considered a negative prognostic factor in several tumor types. The role of ENPP3 in normal non-immune tissues and tumors is less clear although recent findings have described a possible activity in cell proliferation and motility^13^, in embryo implantation^14^, endometrial function^15^ and ovarian regulation^14,16^. It has been shown that some epithelial tissues express high levels of ENPP3 and this expression is maintained also upon malignant transformation and can be detected in renal and biliary carcinomas^17^. For this reason, ENPP3-directed antibodies have been developed and antibody–drug conjugates (ADCs) have shown anti-tumor efficacy in clinical trials for advanced renal cell carcinoma^18–20^. In the same pathology, dual targeting of ENPP3 and SIRPα via bispecific antibodies demonstrated anti-tumor efficacy by promoting tumor killing by macrophages^21^. Finally, SRSF1, by regulating ENPP3 splicing, has been implicated in inflammation through BRD4 de-glycosylation and NF-κB signaling^22^. These findings, together with the increasing attention caught by CD39 and CD73, which regulate extracellular adenosine signaling and strongly contribute to the immunosuppressive tumor microenvironment ^23^, clearly support the notion that ENPP3 is not merely a lineage marker, but it could play a role in tumor biology and immune escape. This activity could stem from its capability to deplete the pro-inflammatory mediator ATP, by producing adenosine, but also via still unknown functions ^8^.

In our work, we provide first evidence that, contrarily to ENPP1, ENPP3 is associated with better prognosis in breast cancer and that it is up-regulated in HER2-positive breast cancer cells. Via immunohistochemical analysis of breast cancer patients and *in vitro* experiments, we show that HER2, together with inflammation and hypoxia, favors ENPP3 expression. Finally, we demonstrate that the depletion of ENPP3 promotes breast cancer aggressiveness, by increasing the epithelial-to-mesenchymal features of *in vivo* mammary cancer models and, most of all, the absence of ENPP3 expression correlates with reduced relapse-free survival in breast cancer patients.

## Methods

### Cell culture and treatments

The mouse mammary PyMT41c (41c) and N2C cell lines were established in our laboratory from spontaneous tumors collected from MMTV-PyMT B6 and BALB/c Neu transgenic mice respectively, as previously described^24^. The other transgenic mouse-derived cancer cell lines used were: TUBO^25^, N202 1A^26^, and N202 1E^27^. The HEK293 cell line was purchased from Thermo Fisher Scientific. Cells were cultured in DMEM (Gibco-Thermo Fisher Scientific) supplemented with 10% fetal bovine serum (FBS; Euroclone), 2 mM L-glutamine (cat. no. 56-85-9; Merck), 1 mM sodium pyruvate (cat. no. 13-115E; Lonza), and 1× non-essential amino acid solution (cat. no. M7145, Merck). Cell lines were cultured at 37°C in a fully humidified atmosphere with 5% CO2, and checked for Mycoplasma with a Mycoplasma Detection Kit (Takara) every month. Cells were treated with ATP (cat.no. R0441; Thermo Fisher Scientific), 20 µM IFN-γ (cat. no 130-105-785, Miltenyi Biotec), 20 ng/ml AREG (cat. no 315-36, Peprotech), 50 ng/ml EPG (cat. no 100-51, Peprotech), 50 ng/ml EREG (cat. no 100-04, Peprotech), 20 ng/ml HB-EGF (cat. no 100-47, Peprotech), 20 ng/ml BTC (cat. no 315-21, Peprotech), 50 ng/ml EGF (cat. no AF-100-15, Peprotech), 20 ng/ml TGFα (cat. no. 100-16A, Peprotech) and recombinant mouse anti-rat HER2 antibody (7.16.4; cat. no. BE027, BioCell).

### Enpp3 ectopic expression and mutagenesis

Plasmid encoding for mouse Enpp3-Myc was purchased by Sino Biological (cat. no MG50940) and was mutagenized with the Q5® Site-Directed Mutagenesis Kit (cat. No. E0554S, New England Biolabs) according to manufacturer’s protocol. The following For_5’-TCCCACCAAAgCCTTCCCAAA-3’ and Rev_5’-TACATAGCTCTCATATATTTAGAATGG-3’ primers were used to introduce the T205A point mutation. HEK293 cells were seeded in 6-well plates and transfected with 2 µg pCMV3-Enpp3-Myc, pcDNA-GFP and an empty pcDNA 3.1 vector together with lipofectamine 2000 reagent (Thermo Fisher Scientific, cat. 11668027). Stably cell lines were obtained by using 500 µg/ml hygromycin selection.

### Generation of the Enpp3-GFP PyMT41c reporter cell line

PyMT41c cells (10^4 cells/well) were seeded in 96-well plates in 100 ul of complete medium. The day after, cells were transfected with 1-5 ul of Lipofectamine 2000 plus 0.2 ug pSpCas9(BB)-2A-GFP gEnpp3 (Addgene 48138) + 0.2 ug of template PCR. Two gRNAs targeting *Enpp3* were cloned: 5’-CACCGGCCTTATTTTCTGGCTAGAC-3’ and 5’-CACCGCTACGGGAACAATGGATTCC-3’. The template for homologous recombination was obtained by PCR amplification of the GFP of TR455_ERE-rFluc-T2A-GFP-mPGK-Puro plasmid (System Biosciences) with primers containing the flanking regions of mouse *Enpp3* (for 5’-TCACAGCTGGGGCAGGCAGGACAGTTCCTTTTTCCCTCCCCAGAGGGAAGAAAGAGCCTTATTTTCTGGCTAGACAGGTTTACACAGCTACGGGAACAATATGGTGAGCAAGGGCGAGGA-3’; rev 5’-CACACAAAGGATCTTGTATTTCTTGAGACTGTCTTTCTTAATGGGCTCCTCTGTGGCTAATGCTAGCCTGGAATCCTTACTTGTACAGCTCGTCCATG-3’. After transfection, single cells were seeded in 96-well plates and screened by PCR to detect integration of GFP. The entire region of the genome containing GFP was then checked by Sanger sequencing.

### Western blot

Cell pellets were resuspended in lysis buffer (125 mM Tris HCl pH 6.8, 5 % SDS) and total cell proteins were extracted by boiling for 10 minutes at 99 °C. Samples were then clarified by centrifugation at 13000 RPM for 15 minutes upon sonication for 20 seconds. Proteins (30-50 μg) were separated by SDS-PAGE on precast 4%– 12% Bis-Tris NuPAGE gels (Thermo Fisher Scientific) and transferred to PVDF membrane (Merck). Membranes were saturated for 30 minutes in Tris-buffered saline containing 4% BSA and incubated overnight with the following primary antibody: Myc-tag (cat. no. 2278, Cell signaling), ENPP3 (cat. no. 75442, Cell signaling), HER2 (cat. no. 06562, Merck), p-ERK1/2 (cat. no. 9271, Cell signaling), p-HER2 (cat. no. 06229, Merck), AKT (cat. no. 9101, Cell signaling), p-AKT (cat. no. 9271, Cell signaling), Vinculin (cat. no. V9131, Merck) and Actin (cat. no. A1978, Merck). Protein images were acquired by using the Azure biosystem 600 (Aurogene) upon hybridization for 1 hour with appropriate secondary antibody HRP-conjugated (anti-mouse cat. no. NA931 or anti-rabbit cat. no. NA934; Merck) chemiluminescent reaction was obtained through the chemiluminescence HRP substrate (Takara).

### 2’-3’cGAMP transfection and detection

HEK293 cells were stimulated with cGAMP via transfection with a mixing solution made of 3.6 µg/ml cGAMP (cat. no. 35573; Cell Signaling Technology) and 3.6 µl lipofectamine 2000 Reagent. After 30 minutes of incubation, the mix was added to recipient cells. To evaluate the level of cGAMP within cells, HEK293 cells treated with cGAMP for 3 and 6 hours were lysed with M-PER mammalian protein extraction reagent (cat.no. 78501; Thermo Fisher Scientific) and analyzed with a cGAMP ELISA kit (cat. no. 501700; Cayman).

### ATP detection assays

HEK293 cells were seeded in 96-well plates at 1x10^4^ cells/well. After 24 hours, the medium was harvested and transferred in a white optical 96-well plate and ATP was added for 3 hours. Levels of ATP were analyzed through CellTiter-Glo Luminescent Cell Viability assay, (cat. no. G7571; Promega), according to manufacturer’s instructions. Luminescence quantification was made by using the microplate reader SPARK (Tecan).

### siRNA transfection

To transiently silence HER2, cells were transfected with a reverse transfection protocol using Lipofectamine RNAiMAX transfection reagent (Thermo Fisher Scientific) with siRNA specific for HER2 (cat. no. SI0063882; Qiagen) and negative control siRNA (cat. no. 3510999, Qiagen) as control of transfection.

### RNA extraction and real-time PCR

Total RNA was isolated from cell pellets with miRNeasy Mini Columns (Qiagen) and 1 μg was reverse-transcribed using the High Capacity cDNA Reverse Transcription Kit (Thermo Fisher Scientific). Real-time PCR to analyze *Enpp3* expression was carried out with TaqMan Fast Universal PCR Master Mix (Thermo Fisher Scientific) by using the Taqman probe Mm01193723_m1 (cat. no. 4331182). Samples were analyzed using QuantStudio 3 software (Thermo Fisher Scientific), and transcript levels of technical duplicates were quantified using the Delta-Ct method and normalized to *Gapdh* (Mm99999915_g1) expression.

### Lentiviral transduction

To produce lentiviral particles, HEK293 cells were seeded in 10 mm petri dishes (5 x10^6^ cells/well) and transfected the day after with viral packaging plasmids (3.75 µg pRSV-REV, 9.75 µg pMDLg/pRRE and 5.25 µg pMD2-VSV-G) and 15 µg of the plasmid of interest. The next day the medium was replaced with fresh one and after further 24 hours passed through a 0.45 mm filter and added to recipient PyMt41c cells, seeded the day before in a 12 well plate (5x10^4^ cells/well). For *Enpp3* knockout, two gRNAs (#2 TTGTTGGTGATTGTATCGCT; #3 CGCTGTGACTCGGGATGCAC) were cloned in LentiCRISPRv2GFP (Addgene, cat. no. 82416) according to Ran et al protocol^28^, but using the BsmBI enzyme, instead of BbsI. Transduction efficiency was determined by GFP expression.

### *In vivo* experiments

Animal experiments were approved by the Ethics Committee for Animal Experimentation of the Fondazione IRCCS Istituto Nazionale dei Tumori of Milan and by the Italian Ministry of Health (authorization #974/2023-PR). C57BL/6^W-sh/W-sh^ (Wsh) mice were purchased from the Jackson Laboratory, and maintained under pathogen-free conditions at the animal facility of Fondazione IRCCS Istituto Nazionale dei Tumori (Milan, Italy). Tumor engraftment experiment was made by injecting female Wsh mice with 1x10^5^ PyMT41c cells control or knockout for *Enpp3* and monitored twice a week for tumor growth.

### Patient cohort, follow-up and immunohistochemistry

Authorization for employment of human FFPE was obtained by the Institutional Ethical Committee (INT n° 12/2017). Breast cancer patients of INT cohort were classified as HER2 (n = 100) if tumors were ER-negative, PGR-negative and HER2 was 2+ amplified or 3+. Patients were classified as triple negative (n = 25) if tumors were ER-negative, PGR-negative and HER2 was not 2+ amplified or 3+. Formalin-fixed, paraffin-embedded (FFPE) breast cancer tissue sections (4 μm) from consecutive patients were deparaffinized by the IHC internal facility of INT, rehydrated, and subjected to heat-induced antigen retrieval in 10 mM citrate buffer (pH 6.0) for 15 min. Sections were incubated with a rabbit polyclonal anti-ENPP3 antibody (HPA043772, Sigma-Aldrich), followed by HRP-based detection with DAB as chromogen and hematoxylin counterstaining. A pathologist blindly determined the percentage of ENPP3 positivity. A few cases were excluded due to technical issues during the staining or absence of follow-up.

### Graphs, *in silico* and statistical analysis

Graphs were generated with GraphPad Prism 11.0. Kaplan-Maier from publicly available data were generated with the https://kmplot.com/analysis/index.php?p=home tool^29^. Statistical analysis was performed with GraphPad Prism 11.0. The statistical test is described in each figure legend.

## Results

### ENPP3 is associated with better prognosis in breast cancer despite its potential immunosuppressive activity

ENPP3 is an ectonucleotidase shown to degrade different nucleotides with variable efficiency^1,2^. In particular, it has been shown to regulate mast cell activation by depleting ATP^8^, and to degrade cGAMP, hence preventing STING pathway activation^10^. To investigate its activity, we over-expressed *Enpp3* cDNA and generated mutant sequences by mutagenizing the T205 residue, which has been shown to be essential for both ATPase and cGAMPase activity^1,10,30^. We confirmed the ectopic expression of the wild type and 2 out 3 mutant sequences (Figure 1A) finding similar levels of proteins. One vector failed to express Enpp3 likely to unsuccessful cloning or plasmid recombination during the mutagenizing process (Figure 1A). Transfected HEK293 cells were then tested *in vitro* confirming that Enpp3 reduces the levels of cGAMP (Figure 1B) and of ATP (Figure 1C). As expected, only wild type Enpp3 maintained the ATPase activity, while mutated Enpp3 T205A failed to hydrolyze ATP (Figure 1C).

**Figure 1:**
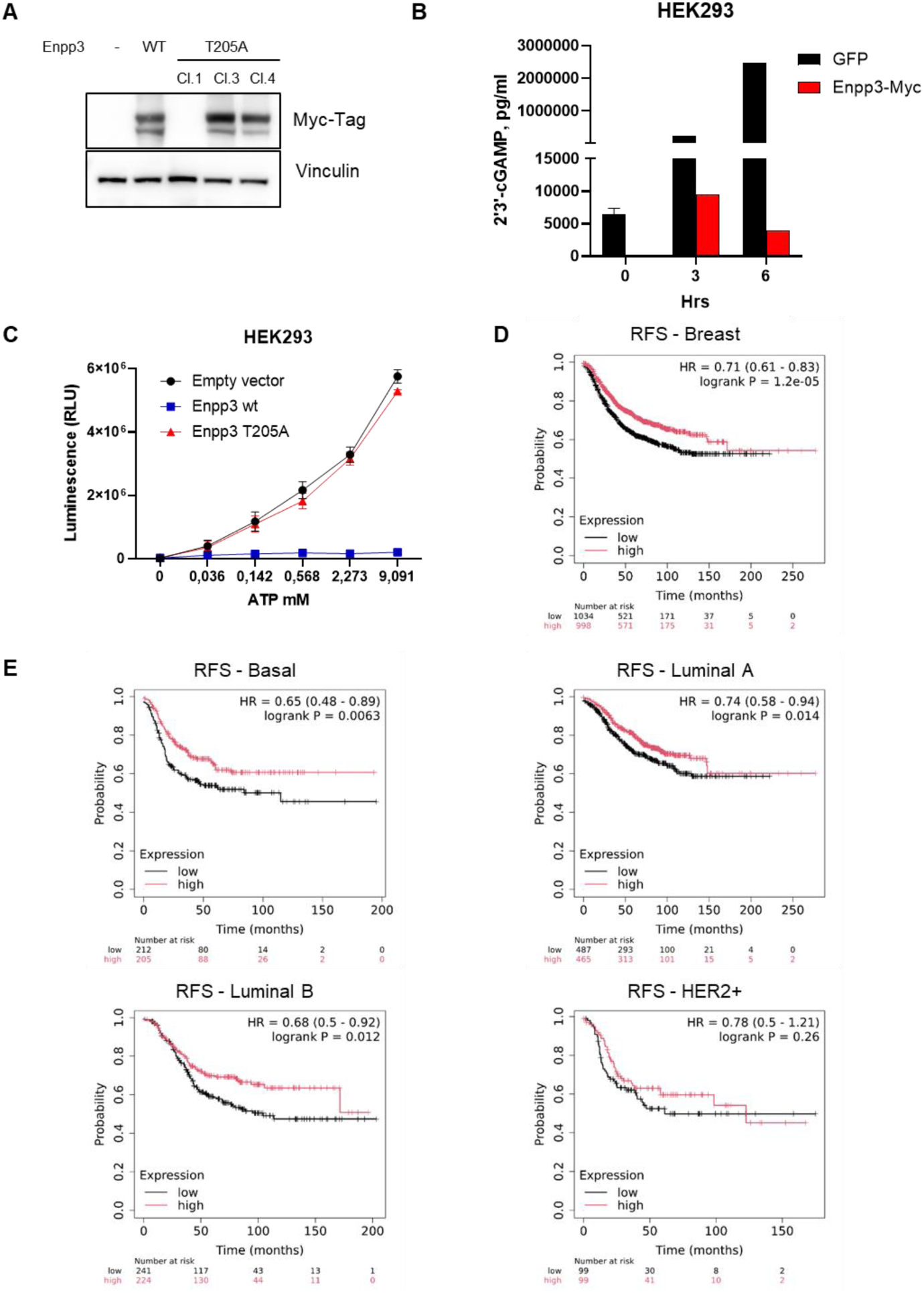
ENPP3 is endowed with ATPase and cGAMPase activity and is associated with better prognosis in breast cancer. (A) Western blot to evaluate the ectopic expression of Enpp3 in HEK293 cells transfected with an empty vector (-), with Myc-tagged wild type (WT) Enpp3 cDNA and with three different Enpp3 T205 mutant sequences (Cl 1, Cl 3 and Cl 4). Ectopic expression was detected with an anti-Myc-tag antibody. Vinculin is shown as a loading control. (B) Levels of cGAMP were evaluated through ELISA in the media of HEK293 cells overexpressing GFP or Enpp3-Myc upon treatment with cGAMP for 3 and 6 hours (n=3). (C) Levels of ATP were measured in the media of HEK293 cells overexpressing wild type Enpp3, T205A Cl3 mutated Enpp3 or transfected with the empty vector. Cells were treated for 3 hours with the indicated concentration of ATP, which was measured via luminescence (n=3). (D) Prognostic value of *ENPP3* in terms of relapse-free survival in (D) all breast cancer patients, and in the (E) Basal, Luminal A, Luminal B and HER2-enriched breast cancer patients^29^.

ENPP3 localizes in the outer region of the plasmatic membrane to which it is anchored thanks to a short transmembrane sequence and from which it can be shed and released in the extracellular space. By depleting ATP and cGAMP, ENPP3 displays a potential immunosuppressive activity, which shares with its closely related homolog ENPP1. Accordingly, ENPP1 expression often correlates with worse prognosis in many cancer types^11,12,31^. Nevertheless, despite structural and functional similarities with ENPP1, *in silico* analysis reveals that ENPP3 is associated with better prognosis in breast cancer (Figure 1D). Its expression is associated with increased relapse-free survival in any breast cancer subtype, not reaching statistical significance only in HER2+ likely due to the limited number of patients in this dataset (Figure 1E). Notably, the same favorable prognostic value is also found in other tumor types (Supplementary Figure 1).

### ENPP3 is more expressed by HER2-positive breast cancer cells

ENPP3 is mainly considered a basophil and mast cell marker^8,32^, although a few works have shown that it can be expressed also by tumors in a tissue-specific manner^21^. In our previous work, we have shown that mast cells enhance the aggressiveness of breast cancer cells by stimulating the activity and expression of estrogen receptor^21^, hence hindering the efficacy of endocrine and anti-HER2 therapy, and by stimulating the stem-like and tumor-initiating properties of cancer cells^22^. Mast cells are known to infiltrate preferentially luminal breast cancer and are associated with worse prognosis, at least in luminal and HER2-positive subtypes ^23^. By analyzing *ENPP3* in different breast cancer subtypes, we found that *ENPP3* is more expressed in HER2-positive breast cancers, both in situ and invasive (Figure 2A). At first, we hence speculated that HER2-positive breast cancers could induce the expression of *ENPP3* by mast cells, but further IHC analysis of breast cancer patients showed that, although mast cells indeed express ENPP3 (Figure 2B), unexpectedly, also breast cancer cells are positive in ENPP3 immunohistochemistry (Figure 2C-E). In these cells, it was appreciable a cytoplasmic staining with membrane reinforcement in agreement with the expected transmembrane localization of this ectoenzyme. Moreover, there was a presence of reinforcement at the level of pseudolumina formed by neoplastic cells and in some spots at the cell junctions (Figure 2C-E). In agreement with *in silico* analyses (Figure 2A), we also observed that in situ tumors (figure 2C-D) expressed higher levels compared to invasive breast cancer (Figure 2E). Interestingly, a previous work has shown that ENPP3-related homolog ENPP1 is expressed in a HER2delta16-dependent manner^24^ suggesting that activated HER2 could stimulate the expression of both ENPP members. Finally, to corroborate the *in silico* analysis, we compared HER2-positive and triple negative breast cancers from our cohort, finding that indeed HER2-positive cells significantly expressed more ENPP3 compared to triple negative/basal breast cancer (Figure 2F).

**Figure 2.**
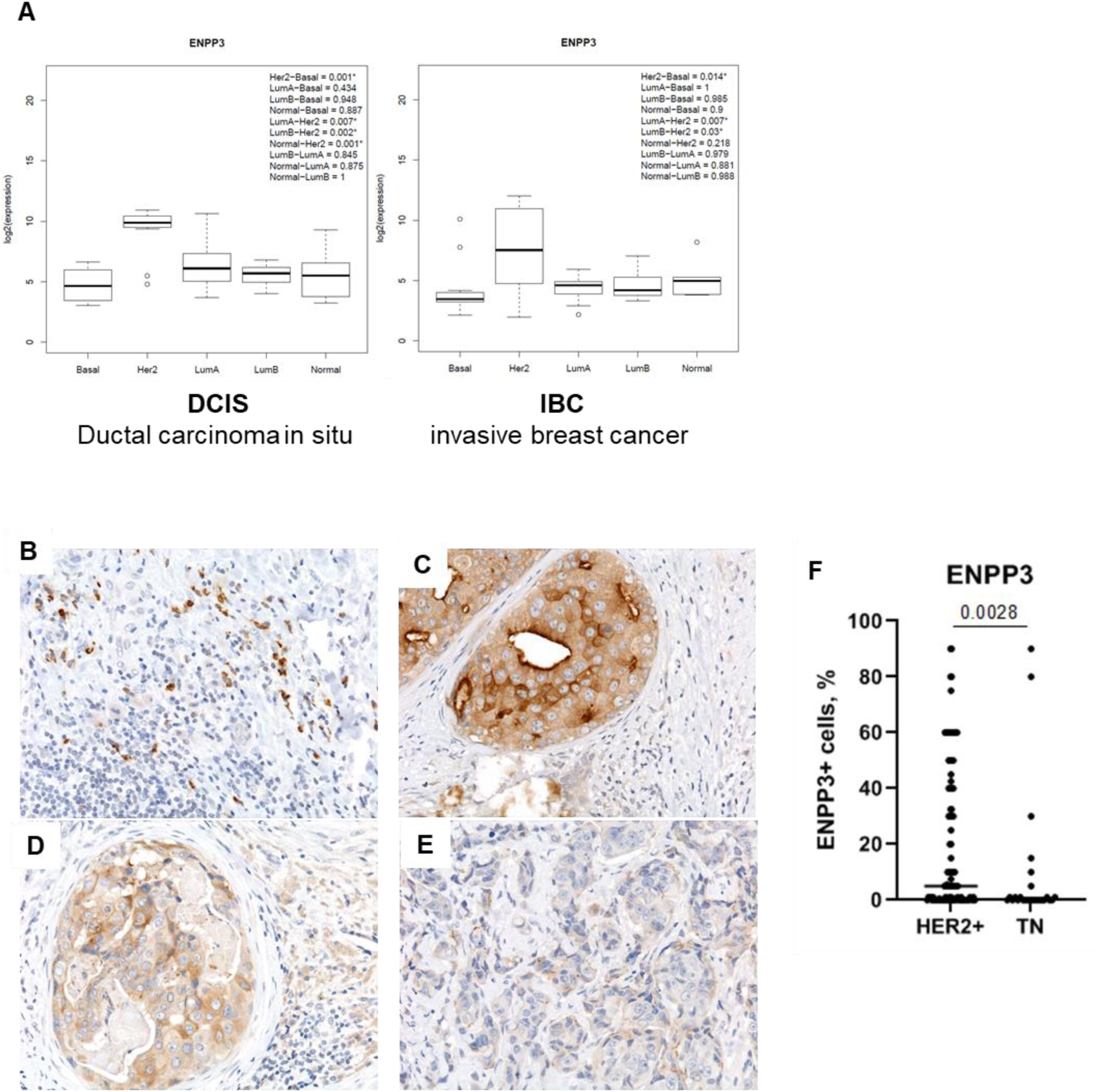
HER2-positive breast cancer cells express higher levels of ENPP3. (A) I*n silico* analysis of *ENPP3* expression in ductal carcinoma in situ (DCIS) and invasive breast cancer (IBC). Immunohistochemical analysis of breast cancer patient FFPE showing (B) immune cells, (C-D) ductal carcinoma in situ and (E) invasive breast cancer patients. (F) Evaluation of percentage of ENPP3 positivity in HER2-positive (n = 100) and triple negative (n=25) breast tumors, Mann Whitney test.

### HER2 is associated with ENPP3 expression *in vitro*

Having shown that HER2-positive breast cancer cells express higher levels of *ENPP3* (Figure 2), we checked its expression levels in mouse mammary cancer models. Among the cell lines tested, Enpp3 was detectable at a protein level in 3 cell lines (N2C, N202 1A and TUBO), all expressing HER2 (Figure 3A). On the contrary, 41c and N202 1E cells, both negative for HER2, do not express Enpp3. Interestingly, N202 1E is a cell line derived from the same transgenic mice as N202 1A, but that lost the expression of transgenic HER2. To further test whether HER2 is responsible for Enpp3 expression, we silenced HER2 in TUBO cells, finding that this was sufficient to reduce Enpp3 expression (Figure 3B) and the same effect was obtained with an antibody against HER2 (Figure 3C). Since ENPP3 prevents excessive mast cell inflammation, and hence it is a regulator of cell inflammation, we checked whether IFN stimulates its expression in cancer cells, but it did not result in the expected effect (Fig. 3B). We then asked whether HER2 activation could stimulate Enpp3 expression. Since no ligand is known to directly activate HER2, we treated N2C cells with a number of EGFR ligands, which, by activating EGFR, ultimately resulted in HER2 activation. The vast majority of ligands increased *Enpp3* transcript (Figure 3D), together with the protein (Figure 3E), confirming that indeed the HER2/EGFR downstream pathway promotes the expression of ENPP3.

**Figure 3.**
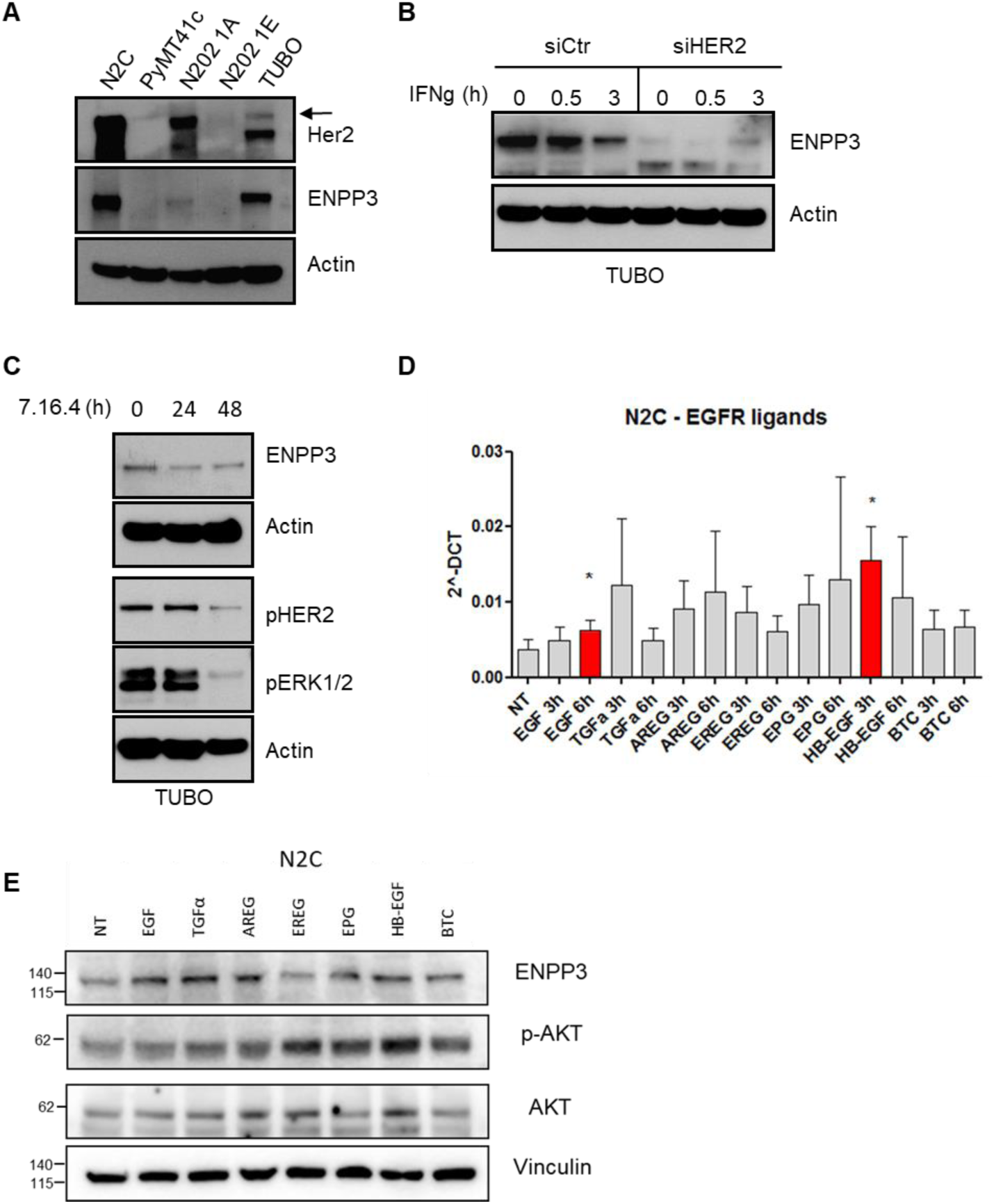
The HER2/EGFR axis stimulates ENPP3 expression. (A) Western blot to evaluate the levels of Enpp3 expression in mouse mammary cancer cell lines (N2C, PyMT41c, N202 1A, N202 1E and TUBO) over-expressing or not HER2. Actin is shown as loading control. (B) Enpp3 levels in TUBO cells transfected with a non-targeting siRNA (siCtr) or with a HER2-specific siRNA (siHER2). Cells were stimulated with IFNg for the indicated times. Actin is shown as a loading control. (C) TUBO cells were treated with the anti-HER2 7.16.4 for 24 and 48 hours. Cells were harvested and analyzed by western blot to evaluate the levels of Enpp3 and the activation of HER2 and ERK1/2. Actin is shown as a loading control. (D) Real Time PCR was performed to evaluate the levels of Enpp3 in N2C cells stimulated with EGFR ligands (epidermal growth factor (EGF), transforming growth factor alpha (TGFα), amphiregulin (AREG), epiregulin (EREG), epigen (EPG), heparin binding-epidermal growth factor (HB-EGF) and betacellulin (BTC) for the indicated times (n=3, One-way ANOVA). (E) The same cells as in (D) were analyzed in western blot, upon 24 hours of treatment, to evaluate the protein levels of Enpp3, together with AKT and its activating phosphorylation. Vinculin is shown as a loading control.

### ENPP3 expression is stimulated by pathways triggered by extracellular signals

Little information is still available regarding the mechanisms by which ENPP3 is expressed. We hence engineered the MMTV-PyMT transgenic mice-derived 41c cell line and integrated a GFP tag under the control of the *Enpp3* promoter with CRISPR/Cas9 technology in order to identify which stimuli could transactive Enpp3 gene expression (Figure 4A). Although 41c cells do not express detectable levels of *Enpp3*, we observed the presence of a few GFP-positive cells in the mass culture of this newly generated reporter cell line. We hence decided to sort cells and perform gene expression analysis by comparing GFP-positive and -negative cells to identify pathways associated or that could stimulate *Enpp3* expression. Although only minor modifications in gene expression were identified, a number of genes were modulated between GFP-positive and -negative populations (Figure 4B). Moreover, pathway analysis showed that NF-kB, IL-6 and hypoxia pathway up-regulation is associated with activation of the *Enpp3* promoter (Figure 4C-D). All these pathways are triggered by extracellular stimuli, together with KRAS signaling, which was inversely correlated with *Enpp3* expression.

**Figure 4.**
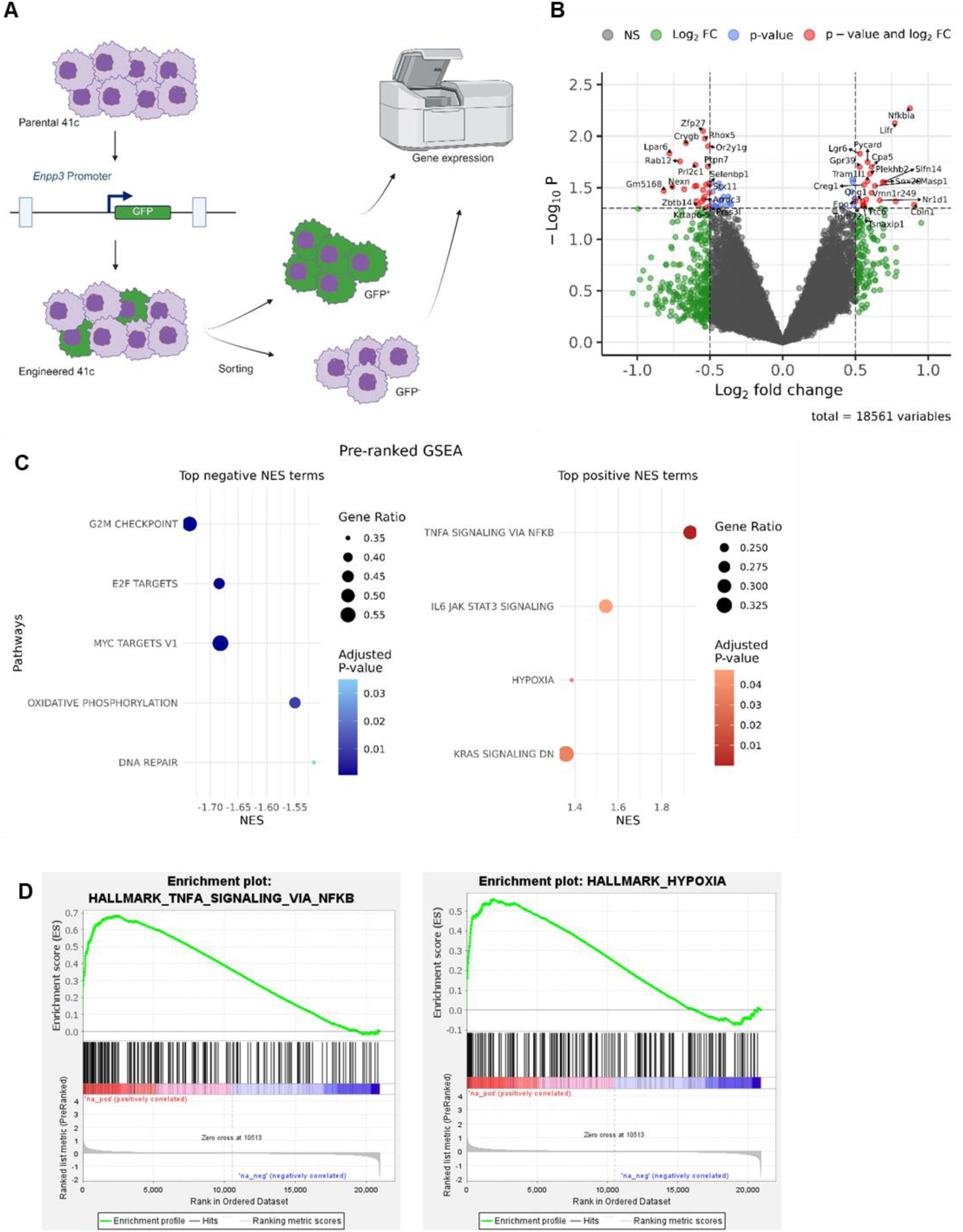
ENPP3 expression is associated with activation by extracellular stimulation. (A) PyMT41c cells were engineered by CRISPR/Cas9 to insert a GFP under the control of the *Enpp3* promoter. Cells were sorted to separate the few GFP-positive from GFP-negative cells; then both populations were profiled by gene expression. (B) Volcano plot showing modulated genes between GFP-positive and GFP-negative cells (n=3, p-values, and not adj p-values, were used for the graph), single genes are shown in Supplementary Table 1. (C) Pathway analysis of the gene expression profiling and (D) enrichment plot with pathways significantly perturbed in the two populations are shown.

### ENPP3 knockout is associated with an EMT-like phenotype and with reduced IFN signaling

We have previously shown that 41c cells express undetectable levels of Enpp3 *in vitro*, but a few cells in the mass culture are likely to express *Enpp3* transcript. We then injected 41c cells *in vivo* finding that, contrarily to *in vitro*, Enpp3 protein became widely expressed upon engraftment (Fig. 5A). Since mast cells are both enriched in the TME and able to express and secrete ENPP3, to avoid any host confounding factor, 41c cells were grown in Wsh mice, which are characterized by the absence of mast cells due to spontaneous inversion of a regulatory region within the cKit promoter ^33^. Moreover, we engineered the 41c cells to knockout *Enpp3*, finding that knockout condition produced significantly larger tumors compared to controls in Wsh mice (Fig. 5B). *Ex vivo* analysis of these tumors further confirmed that detected Enpp3 was from mammary cancer cells (Fig. 5A). Moreover, *ex vivo* analysis highlighted also that Enpp3 knockout tumors were less infiltrated by immune cells and displayed a much more EMT-like phenotype (Fig.5C). Gene profiling was performed on ex vivo wild type and Enpp3 knockout tumors, which confirmed that EMT is associated with *Enpp3* knockout tumors, which, on the contrary, are characterized by reduced response to interferon pathways (Fig. 5D). We finally asked whether ENPP3 is truly associated with better prognosis in breast cancer patients. Follow up of patients belonging to the small cohort shown in Figure 2F showed that HER2-positive patients characterized by tumors expressing ENPP3 are less likely to face relapse (Fig. 5E), hence confirming that, in HER2-positive breast cancer, ENPP3 is indeed associated with better prognosis.

**Figure 5.**
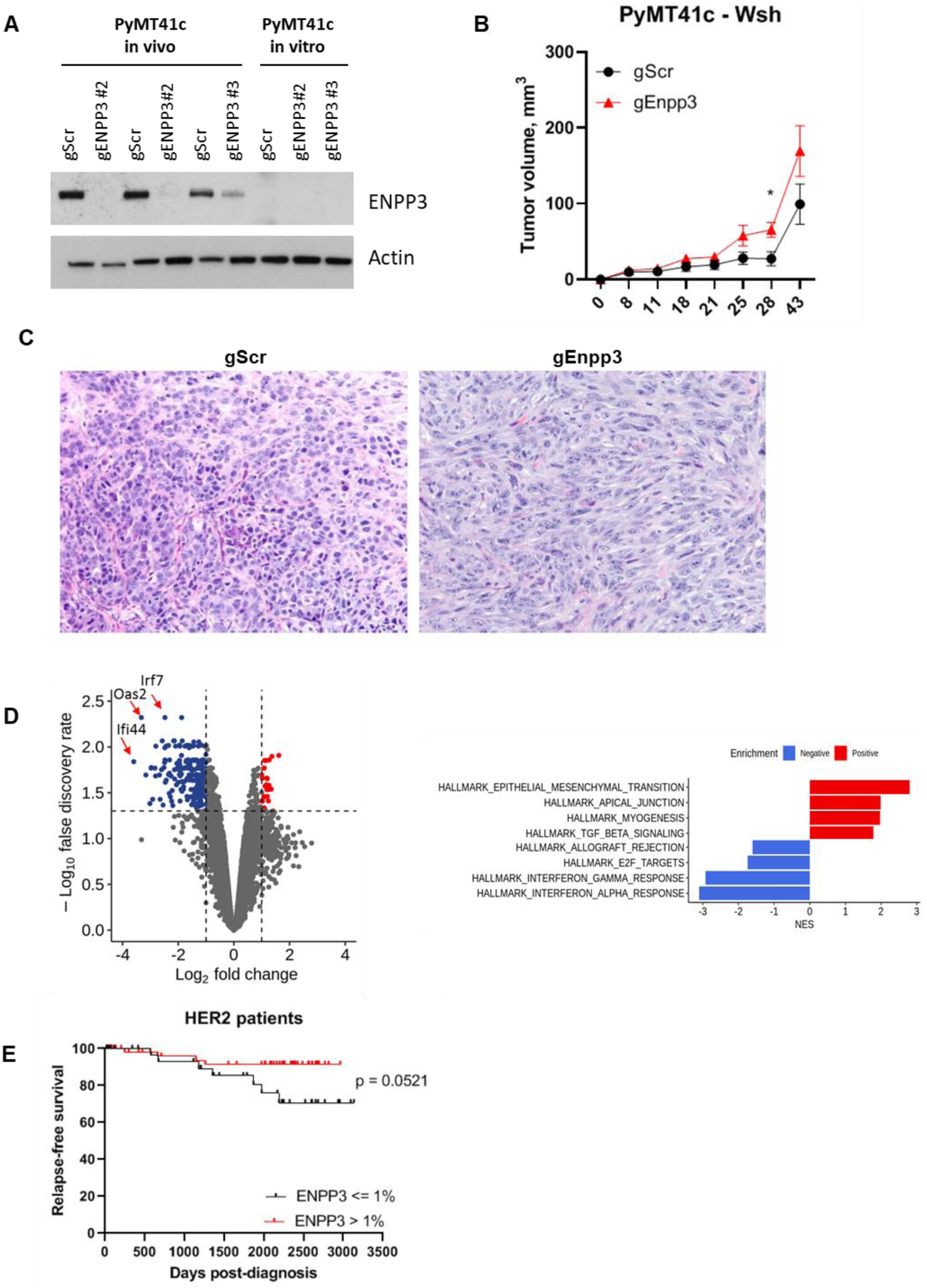
ENPP3 depletion is associated with epithelial-to-mesenchymal breast cancer phenotype and increased relapse. (A) PyMT41c cells were engineered to knockout *Enpp3* with specific gRNAs or with a control non-targeting gRNA (gScr). Western blot was performed to evaluate the levels of Enpp3 in both *ex vivo* tumors injected in Wsh mice and of the same PyMT41c cells cultured *in vitro*. Actin is shown as a loading control. (B) Tumor size of *Enpp3* knockout and control PyMT41c cells injected in Wsh mice. (C) Hematoxylin and eosin staining of representative control and knockout *ex vivo* PyMT41c tumors. (D) Volcano plot of differentially expressed genes between five control and five *Enpp3* knockout *ex vivo* PyMT41c tumors (left) and pathway analysis (right). (E) Relapse free survival of HER2-positive patients according to ENPP3 expression (ENPP3 < 1% n = 36, ENPP3 ≥ 1% n = 59, p-value Log rank test).

## Discussion

In our work, we demonstrate for the first time that ENPP3 is expressed by breast cancer cells, showing that HER2 positivity and stimulation correlate with its up-regulation. We also provide evidence that ENPP3, differently from other ectonucleotidases such as ENPP1, CD73 and CD39, is associated with reduced tumor aggressiveness in mouse models, and with increased relapse-free survival in patients. Our findings suggest that ENPP3 expression prevents the epithelial-to-mesenchymal phenotype, which is a well-known marker of tumor aggressiveness.

The growing consciousness that the tumor microenvironment plays a central role in cancer development, metastasis dissemination and response to therapy has attracted increasing attention towards players that can perturb stromal rearrangement and immune infiltration. Several studies have shown that proteins such as CD73 and CD39 reduce the levels of ATP ^23^, which normally triggers an anti-tumor effect by activating dendritic cells, macrophages and lymphocytes ^34^, and promote the accumulation of adenosine. The latter promotes an immunosuppressive environment, rich of T regulatory cells (Tregs), M2-like macrophages and anergic CD8 T cells ^23^. ENPP1 is a closely related homolog of ENPP3 and shares some activities and substrates with it, being able to hydrolyze ATP and, especially, cGAMP. This molecule is crucial for the activation of the interferon pathways through the activation of the STING cascade. By efficiently depleting cGAMP ^35^, ENPP1 promotes tumor development and resistance to therapy ^36^, immunoevasion and metastasis dissemination^11^, and it is considered a good candidate for targeted therapy ^37,38^ to favor an anti-tumor immune response. Accordingly, it has been shown that cGAMP analogs resistant to ENPP1 hydrolysis stimulate interferon and could be used as adjuvants in vaccine-based anti-tumor treatments ^39^. Not surprisingly, ENPP1 is considered a marker of worse prognosis ^40^. On the contrary, we show here that ENPP3, which has recently been demonstrated to play redundant cGAMP-hydrolysing activity with ENPP1 ^10^, is associated with better prognosis and correlates with increased relapse-free survival. Moreover, our *in vivo* results suggest that *Enpp3* knockout reduces immune infiltration and favors the development of more aggressive tumors.

Hence, ENPP3 could play both anti-tumor cell-intrinsic and pro-tumor cell-extrinsic effects (by contributing to an immunosuppressive microenvironment), and, in our model, the cell-intrinsic effect is apparently prevalent. In fact, our findings support the intriguing possibility that ENPP3 could reduce the aggressiveness of tumors by favoring an epithelial phenotype. Its knockout favored EMT eventually resulting in a counterintuitive immunosuppressed microenvironment. This could be possible via several mechanisms as EMT can hinder the immune-mediated control of tumors by promoting MHC down-regulation and increasing the expression of immune checkpoints and the recruitment of suppressive immune populations ^41–44^. In this setting, ENPP3 would have a different role compared to the one described in mast cells^8^. However, also in our work we provide evidence that TNF-mediated inflammation and other cytokines, via NF-kB, could be associated with ENPP3 expression. Importantly, ENPP3 is extensively investigated in mast cells and basophils, but our, and others’ ^19^, work have highlighted that it is expressed also in non-immune cell types. This observation implies that deconvoluting mast cell infiltration in tumors via expression levels of ENPP3 could be biased by tissue expression and that ENPP3 cannot be considered a *bona fide* marker of mast cells and basophils.

Moreover, for the first time, we describe the capability of HER2 to up-regulate ENPP3 levels, providing a novel mechanism of action which distinguishes ENPP3 from its paralog ENPP1. In fact, although the latter is widely expressed, ENPP3 is more confined to specific tissues both in mice ^10^ and humans ^19^. Since its expression is maintained in transformed cells, ENPP3 has been exploited for antibody-mediated target therapy ^18,20,21,45^ and CAR-T approaches ^46^. Our results support the idea that ENPP3 could be leveraged to target specific tumors, but that its inhibition could be detrimental, being associated with better prognosis. Further work is necessary to understand the biological significance of HER2-mediated regulation, but, since this receptor is responsible for the stimulation of EMT ^47^, while our data support an opposite role for ENPP3, we speculate that ENPP3 could represent a negative feedback or a regulator of HER2/EGFR signaling pathways. Notably, HER2-d16, a constitutively active isoform of HER2, was shown to stimulate the expression of ENPP1, which contributes to cancer immune evasion ^31^. This evidence further strengthens our observation that HER2 is able to promote the expression Enpp members, which could display different activities in the biology of cancer cells.

## Conclusion

With our work, we describe for the first time the expression of ENPP3 in HER2-positive breast cancer cells, providing evidence that ENPP3 is a novel positive prognostic factor. Although mechanistic experiments are still necessary to understand the molecular activity of ENPP3, our work supports the notion that it could be a negative regulator of EMT in breast cancer cells.

## Acknowledgements

The research leading to these results has received funding from AIRC under IG 2020 -ID. 24309 project – P.I. Lecis Daniele. D. Lecis received funds from Italian Ministry of Health “Ricerca Corrente” and 5×1000 Funds– 2013, Italian Ministry of Health.

## References

1 Gorelik A, Randriamihaja A, Illes K, Nagar B. Structural basis for nucleotide recognition by the ectoenzyme CD203c. FEBS J 2018; 285: 2481–2494.

2 Reigada D, Lu W, Zhang X, Friedman C, Pendrak K, McGlinn A et al. Degradation of extracellular ATP by the retinal pigment epithelium. Am J Physiol Cell Physiol 2005; 289: C617–624.

3 Pan C-T, Lin C-C, Lin I-J, Chien K-Y, Lin Y-S, Chang H-H et al. The evolution and structure of snake venom phosphodiesterase (svPDE) highlight its importance in venom actions. Elife 2023; 12: e83966.

4 Dsouza C, Moussa MS, Mikolajewicz N, Komarova SV. Extracellular ATP and its derivatives provide spatiotemporal guidance for bone adaptation to wide spectrum of physical forces. Bone Rep 2022; 17: 101608.

5 Tsai SH, Takeda K. Regulation of allergic inflammation by the ectoenzyme E-NPP3 (CD203c) on basophils and mast cells. Semin Immunopathol 2016; 38: 571–579.

6 Hauswirth AW, Escribano L, Prados A, Nuñez R, Mirkina I, Kneidinger M et al. CD203c is overexpressed on neoplastic mast cells in systemic mastocytosis and is upregulated upon IgE receptor cross-linking. Int J Immunopathol Pharmacol 2008; 21: 797–806.

7 Grootens J, Ungerstedt JS, Wu C, Hamberg Levedahl K, Nilsson G, Dahlin JS. CD203c distinguishes the erythroid and mast cell-basophil differentiation trajectories among human FcεRI+ bone marrow progenitors. Allergy 2020; 75: 211–214.

8 Tsai SH, Kinoshita M, Kusu T, Kayama H, Okumura R, Ikeda K et al. The ectoenzyme E-NPP3 negatively regulates ATP-dependent chronic allergic responses by basophils and mast cells. Immunity 2015; 42: 279–293.

9 Qi Z, Xue Q, Wang H, Cao B, Su Y, Xing Q et al. Serum CD203c+ Extracellular Vesicle Serves as a Novel Diagnostic and Prognostic Biomarker for Succinylated Gelatin Induced Perioperative Hypersensitive Reaction. Front Immunol 2021; 12: 732209.

10 Mardjuki R, Wang S, Carozza J, Zirak B, Subramanyam V, Abhiraman G et al. Identification of the extracellular membrane protein ENPP3 as a major cGAMP hydrolase and innate immune checkpoint. Cell Rep 2024; 43: 114209.

11 Wang S, Böhnert V, Joseph AJ, Sudaryo V, Skariah G, Swinderman JT et al. ENPP1 is an innate immune checkpoint of the anticancer cGAMP-STING pathway in breast cancer. Proc Natl Acad Sci U S A 2023; 120: e2313693120.

12 Carozza JA, Cordova AF, Brown JA, AlSaif Y, Böhnert V, Cao X et al. ENPP1’s regulation of extracellular cGAMP is a ubiquitous mechanism of attenuating STING signaling. Proc Natl Acad Sci U S A 2022; 119: e2119189119.

13 Zhou M, Qi L, Gu Y. GRIA2/ENPP3 Regulates the Proliferation and Migration of Vascular Smooth Muscle Cells in the Restenosis Process Post-PTA in Lower Extremity Arteries. Front Physiol 2021; 12: 712400.

14 Li FX, Yu JJ, Liu Y, Miao XP, Curry TE Jr. Induction of Ectonucleotide Pyrophosphatase/Phosphodiesterase 3 During the Periovulatory Period in the Rat Ovary. ReprodSci 2016; 24: 1033–1040.

15 Chen Q, Xin A, Qu R, Zhang W, Li L, Chen J et al. Expression of ENPP3 in human cyclic endometrium: a novel molecule involved in embryo implantation. Reprod Fertil Dev 2018; 30: 1277–1285.

16 Qin Y, Li Y, Hao Y, Li Y, Kang S. Hypomethylation of the ENPP3 promoter region contributes to the occurrence and development of ovarian endometriosis via the AKT/mTOR/4EBP1 signaling pathway. Biomol Biomed 2023; 24: 848–856.

17 Yano Y, Hayashi Y, Sano K, Nagano H, Nakaji M, Seo Y et al. Expression and localization of ecto-nucleotide pyrophosphatase/phosphodiesterase I-1 (E-NPP1/PC-1) and -3 (E-NPP3/CD203c/PD-Ibeta/B10/gp130(RB13-6)) in inflammatory and neoplastic bile duct diseases. Cancer Lett 2004; 207: 139–147.

18 Thompson JA, Motzer RJ, Molina AM, Choueiri TK, Heath EI, Redman BG et al. Phase I Trials of Anti-ENPP3 Antibody-Drug Conjugates in Advanced Refractory Renal Cell Carcinomas. Clin Cancer Res 2018; 24: 4399–4406.

19 Donate F, Raitano A, Morrison K, An Z, Capo L, Avina H et al. AGS16F Is a Novel Antibody Drug Conjugate Directed against ENPP3 for the Treatment of Renal Cell Carcinoma. ClinCancer Res 2016; 22: 1989–1999.

20 Zhao H, Gulesserian S, Ganesan SK, Ou J, Morrison K, Zeng Z et al. Inhibition of Megakaryocyte Differentiation by Antibody-Drug Conjugates (ADCs) is Mediated by Macropinocytosis: Implications for ADC-induced Thrombocytopenia. Mol Cancer Ther 2017; 16: 1877–1886.

21 Xu L, Wang S, Li D, Yang B, Zhang J, Ran L et al. Dual targeting of ENPP3 and SIRPα with a bispecific antibody enhances macrophage-mediated immunity in renal cell carcinoma. Biochem Biophys Res Commun 2025; 769: 151955.

22 Hu C-K, He L, Huang W-Z, Huang Y, Dai R-X, Chang C et al. Divergent splicing factor SRSF1 signaling promotes inflammation post-CME: the SRSF1/ENPP3 axis acts via inhibition of BRD4 O-GlcNAcylation to enhance NF-κB activation and accelerate heart failure. Theranostics 2025; 15: 6839–6856.

23 Allard B, Longhi MS, Robson SC, Stagg J. The ectonucleotidases CD39 and CD73: Novel checkpoint inhibitor targets. Immunol Rev 2017; 276: 121–144.

24 Majorini MT, Cancila V, Rigoni A, Botti L, Dugo M, Triulzi T et al. Infiltrating mast cell-mediated stimulation of estrogen receptor activity in breast cancer cells promotes the luminal phenotype. Cancer Res 2020; 80: 2311–2324.

25 Rovero S, Amici A, Di Carlo E, Bei R, Nanni P, Quaglino E et al. DNA vaccination against rat her-2/Neu p185 more effectively inhibits carcinogenesis than transplantable carcinomas in transgenic BALB/c mice. J Immunol 2000; 165: 5133–5142.

26 Lollini PL, Nicoletti G, Landuzzi L, De Giovanni C, Rossi I, Di Carlo E et al. Down regulation of major histocompatibility complex class I expression in mammary carcinoma of HER-2/neu transgenic mice. Int J Cancer 1998; 77: 937–941.

27 Nanni P, Nicoletti G, De Giovanni C, Landuzzi L, Di Carlo E, Cavallo F et al. Combined allogeneic tumor cell vaccination and systemic interleukin 12 prevents mammary carcinogenesis in HER-2/neu transgenic mice. J Exp Med 2001; 194: 1195–1205.

28 Ran FA, Hsu PD, Wright J, Agarwala V, Scott DA, Zhang F. Genome engineering using the CRISPR-Cas9 system. NatProtoc 2013; 8: 2281–2308.

29 Posta M, Győrffy B. Pathway-level mutational signatures predict breast cancer outcomes and reveal therapeutic targets. Br J Pharmacol 2025; 182: 5734–5747.

30 Korekane H, Park JY, Matsumoto A, Nakajima K, Takamatsu S, Ohtsubo K, et al. Identification of ectonucleotide pyrophosphatase/phosphodiesterase 3 (ENPP3) as a regulator of N-acetylglucosaminyltransferase GnT-IX (GnT-Vb). JBiolChem 2013; 288: 27912–27926.

31 Attalla SS, Boucher J, Proud H, Taifour T, Zuo D, Sanguin-Gendreau V et al. HER2Δ16 Engages ENPP1 to Promote an Immune-Cold Microenvironment in Breast Cancer. Cancer Immunol Res 2023; 11: 1184– 1202.

32 Marone G, Varricchi G, Loffredo S, Granata F. Mast cells and basophils in inflammatory and tumor angiogenesis and lymphangiogenesis. EurJPharmacol 2016; 778: 146–151.

33 Grimbaldeston MA, Chen CC, Piliponsky AM, Tsai M, Tam SY, Galli SJ. Mast cell-deficient W-sash c-kit mutant Kit W-sh/W-sh mice as a model for investigating mast cell biology in vivo. AmJPathol 2005; 167: 835–848.

34 Di Virgilio F, Sarti AC, Falzoni S, De Marchi E, Adinolfi E. Extracellular ATP and P2 purinergic signalling in the tumour microenvironment. Nat Rev Cancer 2018; 18: 601–618.

35 Kato K, Nishimasu H, Oikawa D, Hirano S, Hirano H, Kasuya G et al. Structural insights into cGAMP degradation by Ecto-nucleotide pyrophosphatase phosphodiesterase 1. Nat Commun 2018; 9: 4424.

36 Carozza JA, Böhnert V, Nguyen KC, Skariah G, Shaw KE, Brown JA et al. Extracellular cGAMP is a cancer cell-produced immunotransmitter involved in radiation-induced anti-cancer immunity. Nat Cancer 2020; 1: 184–196.

37 Pu C, Cui H, Yu H, Cheng X, Zhang M, Qin L et al. Oral ENPP1 inhibitor designed using generative AI as next generation STING modulator for solid tumors. Nat Commun 2025; 16: 4793.

38 Cho Y, Kang M, Ji SH, Jeong HJ, Jung JE, Oh DH et al. Discovery of Orally Bioavailable Phthalazinone Analogues as an ENPP1 Inhibitor for STING-Mediated Cancer Immunotherapy. J Med Chem 2023; 66: 15141–15170.

39 Li L, Yin Q, Kuss P, Maliga Z, Millán JL, Wu H et al. Hydrolysis of 2’3’-cGAMP by ENPP1 and design of nonhydrolyzable analogs. Nat Chem Biol 2014; 10: 1043–1048.

40 Ruiz-Fernández de Córdoba B, Moreno H, Valencia K, Perurena N, Ruedas P, Walle T et al. Tumor ENPP1 (CD203a)/Haptoglobin Axis Exploits Myeloid-Derived Suppressor Cells to Promote Post-Radiotherapy Local Recurrence in Breast Cancer. Cancer Discov 2022; 12: 1356–1377.

41 Imodoye SO, Adedokun KA. EMT-induced immune evasion: connecting the dots from mechanisms to therapy. Clin Exp Med 2023; 23: 4265–4287.

42 Dongre A, Rashidian M, Reinhardt F, Bagnato A, Keckesova Z, Ploegh HL et al. Epithelial-to-Mesenchymal Transition Contributes to Immunosuppression in Breast Carcinomas. Cancer Res 2017; 77: 3982–3989.

43 Dongre A, Rashidian M, Eaton EN, Reinhardt F, Thiru P, Zagorulya M et al. Direct and Indirect Regulators of Epithelial-Mesenchymal Transition-Mediated Immunosuppression in Breast Carcinomas. Cancer Discov 2021; 11: 1286–1305.

44 Wang G, Xu D, Zhang Z, Li X, Shi J, Sun J et al. The pan-cancer landscape of crosstalk between epithelial-mesenchymal transition and immune evasion relevant to prognosis and immunotherapy response. NPJ Precis Oncol 2021; 5: 56.

45 Kollmannsberger C, Choueiri TK, Heng DYC, George S, Jie F, Croitoru R et al. A Randomized Phase II Study of AGS-16C3F Versus Axitinib in Previously Treated Patients with Metastatic Renal Cell Carcinoma. Oncologist 2021; 26: 182–e361.

46 Okada R, Mendoza A, Nousome D, Mathur S, Rodriguez C, Oh J et al. ENPP3 CAR T cells combined with CD206 modulation suppress adrenocortical carcinoma. J Immunother Cancer 2026; 14: e013726.

47 Gupta P, Srivastava SK. HER2 mediated de novo production of TGFβ leads to SNAIL driven epithelial-to-mesenchymal transition and metastasis of breast cancer. Mol Oncol 2014; 8: 1532–1547.

