## Supplemental material for "ENPP3 expressed by HER2-positive breast cancer cells is associated with good prognosis by restraining epithelial-to-mesenchymal phenotype"

### Supplementary Figure 1

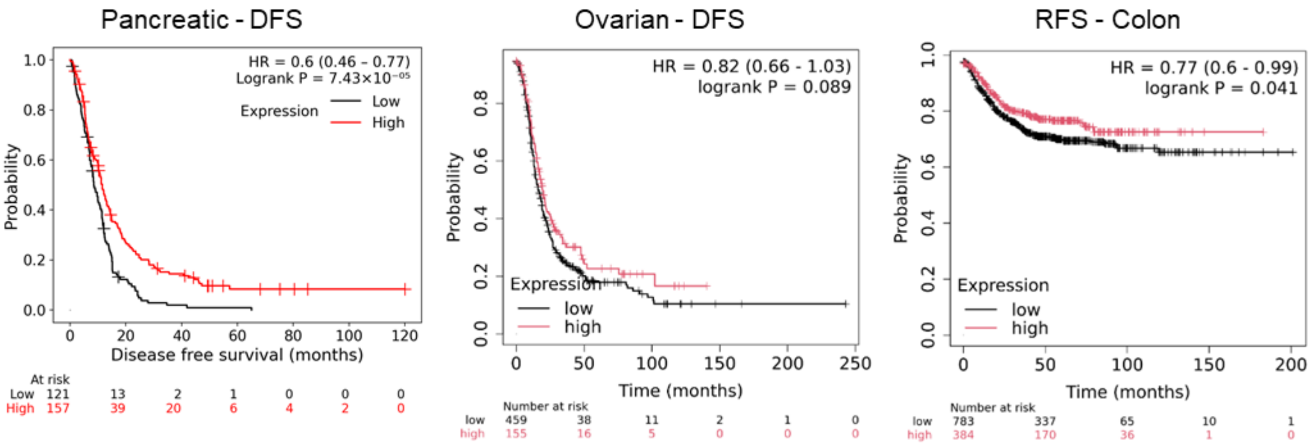

**Supplementary Figure 1. *ENPP3* expression is associated with better prognosis in different tumor types.** Prognostic value of *ENPP3* in terms of relapse-free survival in pancreatic, ovarian and colon cancer patients [29].

**Supplementary Table 1. Genes modulated in association to *Enpp3* promoter activity.**

| ENTREZID | PROBEID | SYMBOL | GENETYPE | GENENAME | logFC | AveExpr | t | P.Value | adj.P.Val | B |
|---|---|---|---|---|---|---|---|---|---|---|
| 18035 | TC1200001768.mm.2 | Nfkbia | protein-coding | nuclear factor of kappa light polypeptide gene enhancer in B cells inhibitor, alpha | 0.875168 | 10.21076 | 4.012179 | 0.005576 | 0.999814 | -4.19155 |
| 16880 | TC1500000045.mm.2 | Lifr | protein-coding | LIF receptor alpha | 0.771533 | 6.037517 | 3.795325 | 0.007312 | 0.999814 | -4.20927 |
| 22689 | TC0700002736.mm.2 | Zfp27 | protein-coding | zinc finger protein 27 | -0.54745 | 6.699748 | -3.6862 | 0.008379 | 0.999814 | -4.21845 |
| 18617 | TC0X00000354.mm.2 | Rhox5 | protein-coding | reproductive homeobox 5 | -0.53519 | 7.605798 | -3.55638 | 0.009912 | 0.999814 | -4.23045 |
| 258463 | TC1100000502.mm.2 | Or2y1g | protein-coding | olfactory receptor family 2 subfamily Y member 1G | -0.51282 | 7.544713 | -3.42711 | 0.011746 | 0.999814 | -4.24309 |
| 329252 | TC0100003119.mm.2 | Lgr6 | protein-coding | leucine-rich repeat-containing G protein-coupled receptor 6 | 0.531033 | 9.537317 | 3.276107 | 0.014366 | 0.999814 | -4.25875 |
| 67168 | TC1400001032.mm.2 | Lpar6 | protein-coding | lysophosphatidic acid receptor 6 | -0.77848 | 8.236244 | -3.23305 | 0.015257 | 0.999814 | -4.26371 |
| 66824 | TC0700004351.mm.2 | Pycard | protein-coding | PYD and CARD domain containing | 0.584148 | 8.073488 | 3.117414 | 0.01782 | 0.999814 | -4.27636 |
| 19328 | TC1700002434.mm.2 | Rab12 | protein-coding | RAB12, member RAS oncogene family | -0.70647 | 7.328458 | -3.11813 | 0.017832 | 0.999814 | -4.27652 |
| 320139 | TC0100001244.mm.2 | Ptpn7 | protein-coding | protein tyrosine phosphatase, non-receptor type 7 | -0.51441 | 6.818243 | -3.0676 | 0.019071 | 0.999814 | -4.28207 |
| 71111 | TC0100001112.mm.2 | Gpr39 | protein-coding | G protein-coupled receptor 39 | 0.52876 | 7.902023 | 3.050486 | 0.019525 | 0.999814 | -4.28407 |
| 226971 | TC0100000233.mm.2 | Plekhb2 | protein-coding | pleckstrin homology domain containing, family B (evectins) member 2 | 0.6004 | 8.754342 | 2.926953 | 0.023203 | 0.999814 | -4.29919 |
| 93670 | TC1100001457.mm.2 | Tac4 | protein-coding | tachykinin 4 | 0.47879 | 6.857451 | 2.859814 | 0.025436 | 0.999814 | -4.30734 |
| 20341 | TC0300000880.mm.2 | Selenbp1 | protein-coding | selenium binding protein 1 | -0.50699 | 6.856842 | -2.78678 | 0.028205 | 0.999814 | -4.31684 |
| 74478 | TSUnmapped00000020.mm.2 | Snx29 | protein-coding | sorting nexin 29 | 0.697768 | 7.16352 | 2.777533 | 0.028594 | 0.999814 | -4.31814 |
| 433375 | TC0100001526.mm.2 | Creg1 | protein-coding | cellular repressor of E1A-stimulated genes 1 | 0.558939 | 9.411793 | 2.74624 | 0.029883 | 0.999814 | -4.32225 |
| 69159 | TC1500002194.mm.2 | Rhebl1 | protein-coding | Ras homolog enriched in brain like 1 | -0.59131 | 6.784197 | -2.72801 | 0.030662 | 0.999814 | -4.32467 |
| 272322 | TC0600001738.mm.2 | Bmal2 | protein-coding | basic helix-loop-helix ARNT like 2 | -0.6046 | 6.584555 | -2.72577 | 0.030759 | 0.999814 | -4.32496 |
| 236915 | TC0X00002660.mm.2 | Arhgef9 | protein-coding | CDC42 guanine nucleotide exchange factor 9 | -0.42867 | 8.288559 | -2.72233 | 0.030861 | 0.999814 | -4.32521 |
| 17174 | TC1600001395.mm.2 | Masp1 | protein-coding | MBL associated serine protease 1 | 0.634627 | 8.026244 | 2.720289 | 0.030998 | 0.999814 | -4.32569 |
| 13607 | TC0X00000941.mm.2 | Eda | protein-coding | ectodysplasin-A | -0.54841 | 6.270684 | -2.68269 | 0.032691 | 0.999814 | -4.33074 |
| 11950 | TC0300002681.mm.2 | Atp5pb | protein-coding | ATP synthase peripheral stalk-membrane subunit b | -0.48157 | 9.623632 | -2.6475 | 0.034314 | 0.999814 | -4.33533 |
| 21886 | TC1000000904.mm.2 | Tle2 | protein-coding | transducin-like enhancer of split 2 | -0.37237 | 7.71163 | -2.61586 | 0.035894 | 0.999814 | -4.33969 |
| 22652 | TC0700003399.mm.2 | Mkrn3 | protein-coding | makorin, ring finger protein, 3 | -0.40622 | 6.567247 | -2.6029 | 0.036563 | 0.999814 | -4.34149 |
| 80885 | TC0500003243.mm.2 | Hcar2 | protein-coding | hydroxycarboxylic acid receptor 2 | -0.53779 | 6.887612 | -2.56462 | 0.038669 | 0.999814 | -4.34704 |
| 269610 | TC0400002023.mm.2 | Chd5 | protein-coding | chromodomain helicase DNA binding protein 5 | -0.42041 | 6.389469 | -2.55676 | 0.039053 | 0.999814 | -4.34797 |
| 14727 | TC1000003213.mm.2 | Lilrb4b | protein-coding | leukocyte immunoglobulin-like receptor, subfamily B, member 4B | 0.494477 | 6.517218 | 2.539728 | 0.040069 | 0.999814 | -4.35056 |
| 328479 | TC1400002750.mm.2 | Gm5089 | ncRNA | predicted gene 5089 | -0.36438 | 7.632702 | -2.52349 | 0.040957 | 0.999814 | -4.3527 |

| | | | | | | | | | | |
|---|---|---|---|---|---|---|---|---|---|---|
| 105171 | TC1300000958.mm.2 | Arrdc3 | protein-coding | arrestin domain containing 3 | -0.5434 | 9.260167 | -2.51851 | 0.041303 | 0.999814 | -4.35359 |
| 230784 | TC0400003674.mm.2 | Sesn2 | protein-coding | sestrin 2 | 0.575165 | 8.199924 | 2.509954 | 0.041813 | 0.999814 | -4.35482 |
| 100040223 | TC0Y00000047.mm.2 | Gm20831 | protein-coding | predicted gene, 20831 | -0.55525 | 6.156253 | -2.49755 | 0.042562 | 0.999814 | -4.3566 |
| 258159 | TC0400001318.mm.2 | Or10ak9 | protein-coding | olfactory receptor family 10 subfamily AK member 9 | -0.35514 | 6.639387 | -2.49565 | 0.042624 | 0.999814 | -4.3567 |
| 70153 | TC1300002098.mm.2 | Qng1 | protein-coding | Q-nucleotide N-glycosylase 1 | 0.547693 | 7.797814 | 2.491059 | 0.04296 | 0.999814 | -4.35754 |
| 217166 | TC1100003688.mm.2 | Nr1d1 | protein-coding | nuclear receptor subfamily 1, group D, member 1 | 0.667116 | 9.968247 | 2.490998 | 0.042964 | 0.999814 | -4.35755 |
| 72149 | TC1100004297.mm.2 | Strada | protein-coding | STE20-related kinase adaptor alpha | -0.38448 | 6.260557 | -2.48808 | 0.04309 | 0.999814 | -4.3578 |
| 17299 | TC1000001544.mm.2 | Mettl1 | protein-coding | methyltransferase 1, tRNA methylguanosine | -0.5933 | 6.67161 | -2.48589 | 0.04328 | 0.999814 | -4.35828 |
| 225443 | TC1800001275.mm.2 | Gm94 | protein-coding | predicted gene 94 | -0.48424 | 6.424748 | -2.47864 | 0.043732 | 0.999814 | -4.35934 |
| 333307 | TC0800002320.mm.2 | Trim75 | protein-coding | tripartite motif-containing 75 | -0.42478 | 6.607875 | -2.46967 | 0.044244 | 0.999814 | -4.36047 |
| 76382 | TC0600000301.mm.2 | 1700012A03Rik | protein-coding | RIKEN cDNA 1700012A03 gene | 0.778064 | 6.734514 | 2.468612 | 0.044366 | 0.999814 | -4.36079 |
| 11416 | TC0300002074.mm.2 | Slc33a1 | protein-coding | solute carrier family 33 (acetyl-CoA transporter), member 1 | -0.46183 | 6.544173 | -2.43551 | 0.046527 | 0.999814 | -4.36564 |
| 13137 | TC0100003035.mm.2 | Cd55b | protein-coding | CD55 molecule, decay accelerating factor for complement B | 0.488496 | 8.754173 | 2.431647 | 0.046786 | 0.999814 | -4.36621 |
| 100040233 | TC0600002156.mm.2 | Prss3l | protein-coding | serine protease 3 like | -0.50262 | 6.133735 | -2.43067 | 0.046852 | 0.999814 | -4.36635 |
| 434246 | TC0700001874.mm.2 | Trim72 | protein-coding | tripartite motif-containing 72 | 0.550901 | 7.309936 | 2.423573 | 0.047332 | 0.999814 | -4.3674 |
| 12017 | TC0400002454.mm.2 | Bag1 | protein-coding | BCL2-associated athanogene 1 | -0.45521 | 9.727606 | -2.41883 | 0.0476 | 0.999814 | -4.36794 |
| 22666 | TC1700001110.mm.2 | Zbtb14 | protein-coding | zinc finger and BTB domain containing 14 | -0.58109 | 7.763127 | -2.41825 | 0.047696 | 0.999814 | -4.36818 |
| 14697 | TC0900001012.mm.2 | Gnb5 | protein-coding | guanine nucleotide binding protein (G protein), beta 5 | 0.412473 | 7.397573 | 2.412636 | 0.048027 | 0.999814 | -4.36886 |
| 70853 | TC0100000257.mm.2 | Vwa3b | protein-coding | von Willebrand factor A domain containing 3B | -0.45112 | 7.408719 | -2.40196 | 0.048828 | 0.999814 | -4.3706 |
| 68484 | TC1600002026.mm.2 | Krtap6-5 | protein-coding | keratin associated protein 6-5 | -0.52731 | 7.727527 | -2.38998 | 0.049677 | 0.999814 | -4.37239 |

Supplementary Table 2

| **SYMBOL** | **transcript_cluster_id** | **ENTREZID** | **GENENAME** | **logFC** | **FC** | **AveExpr** | **t** | **P.Value** | **adj.P.Val** |
|---|---|---|---|---|---|---|---|---|---|
| Rbp7 | TC0400004041.mm.2 | 63954 | retinol binding protein 7, cellular | -3.325620016 | -10.02562322 | 8.908605712 | -16.28928524 | 5.93E-07 | 0.004776196 |
| Egln3 | TC1200001742.mm.2 | 112407 | egl-9 family hypoxia-inducible factor 3 | -1.877421061 | -3.674176827 | 10.1365546 | -16.14138222 | 6.33E-07 | 0.004776196 |
| Slc15a2 | TC1600001582.mm.2 | 57738 | solute carrier family 15 (H+/peptide transporter), member 2 | -2.476967118 | -5.567258677 | 6.901995655 | -15.968022 | 6.83E-07 | 0.004776196 |
| Insl6 | TC1900001340.mm.2 | 27356 | insulin-like 6 | -2.138478937 | -4.402975867 | 8.811475908 | -14.07603095 | 1.65E-06 | 0.0085795 |
| Plekhb1 | TC0700003823.mm.2 | 27276 | pleckstrin homology domain containing, family B (evectins) member 1 | -2.165082543 | -4.484920906 | 8.81191428 | -13.16407283 | 2.64E-06 | 0.0085795 |
| C9 | TC1500000034.mm.2 | 12279 | complement component 9 | -1.855020833 | -3.61756975 | 6.823631817 | -13.16008787 | 2.64E-06 | 0.0085795 |
| Mia | TC0700004639.mm.2 | 12587 | melanoma inhibitory activity | -2.57378744 | -5.953703766 | 9.799907709 | -13.01055236 | 2.86E-06 | 0.0085795 |
| Trim30d | TC0700003946.mm.2 | 209387 | tripartite motif-containing 30D | -1.438072601 | -2.709586306 | 8.700244999 | -12.64898478 | 3.48E-06 | 0.008814774 |
| Trf | TC0900003344.mm.2 | 22041 | transferrin | -1.256865081 | -2.389758911 | 11.91421876 | -12.49902849 | 3.78E-06 | 0.008814774 |
| Ceacam1 | TC0700002635.mm.2 | 26365 | carcinoembryonic antigen-related cell adhesion molecule 1 | -1.948882682 | -3.860754136 | 9.263173384 | -12.11905546 | 4.68E-06 | 0.009539928 |
| Slc44a3 | TC0300002840.mm.2 | 213603 | solute carrier family 44, member 3 | -1.168290584 | -2.247452437 | 8.801520057 | -12.00373202 | 5.00E-06 | 0.009539928 |
| Dhx58 | TC1100003779.mm.2 | 80861 | DEXH (Asp-Glu-X-His) box polypeptide 58 | -1.655704238 | -3.150769549 | 8.752833694 | -11.76408822 | 5.75E-06 | 0.009622607 |
| Deptor | TC1500000392.mm.2 | 97998 | DEP domain containing MTOR-interacting protein | -1.461023851 | -2.753036716 | 8.290455016 | -11.360611 | 7.31E-06 | 0.009622607 |

| | | | | | | | | | |
|---|---|---|---|---|---|---|---|---|---|
| Oas1a | TC0500003181.mm.2 | 246730 | 2'-5' oligoadenylate synthetase 1A | -1.668665713 | -3.179204264 | 8.727665384 | -11.16858906 | 8.22E-06 | 0.009622607 |
| Muc20 | TC1600001514.mm.2 | 224116 | mucin 20 | -1.603635911 | -3.039082651 | 8.698328286 | -11.16730751 | 8.22E-06 | 0.009622607 |
| H2-T23 | TC1700002819.mm.2 | 15040 | histocompatibility 2, T region locus 23 | -1.229239315 | -2.34443343 | 9.718095894 | -11.0176831 | 9.02E-06 | 0.009622607 |
| Atp1b1 | TC0100003422.mm.2 | 11931 | ATPase, Na+/K+ transporting, beta 1 polypeptide | -1.700511062 | -3.250160721 | 8.13569692 | -10.9583195 | 9.36E-06 | 0.009622607 |
| Plppr4 | TC0300002820.mm.2 | 229791 | phospholipid phosphatase related 4 | -1.442043161 | -2.717053854 | 7.478103705 | -10.93457409 | 9.50E-06 | 0.009622607 |
| Csn3 | TC0500000802.mm.2 | 12994 | casein kappa | -4.346261615 | -20.34019514 | 10.15273824 | -10.92996381 | 9.53E-06 | 0.009622607 |
| Iigp1 | TC1800000610.mm.2 | 60440 | interferon inducible GTPase 1 | -2.347235925 | -5.088484081 | 9.820296758 | -10.91281134 | 9.63E-06 | 0.009622607 |
| Gbp10 | TC0500002895.mm.2 | 626578 | guanylate-binding protein 10 | -1.620539384 | -3.074899768 | 9.384330633 | -10.70694686 | 1.10E-05 | 0.009806623 |
| Psmb9 | TC1700001902.mm.2 | 16912 | proteasome (prosome, macropain) subunit, beta type 9 (large multifunctional peptidase 2) | -1.758425565 | -3.383287002 | 10.81085925 | -10.53360664 | 1.23E-05 | 0.009806623 |
| Gm12250 | TC1100000681.mm.2 | 631323 | predicted gene 12250 | -1.940880794 | -3.839399791 | 8.808842636 | -10.5332908 | 1.23E-05 | 0.009806623 |
| Igtp | TC1100004265.mm.2 | 16145 | interferon gamma induced GTPase | -2.803869266 | -6.983107908 | 9.035248305 | -10.47758318 | 1.27E-05 | 0.009806623 |
| Sorbs2 | TC0800000516.mm.2 | 234214 | sorbin and SH3 domain containing 2 | -1.548486604 | -2.925101332 | 7.832594984 | -10.40128721 | 1.34E-05 | 0.009806623 |
| Cxcl9 | TC0500002755.mm.2 | 17329 | chemokine (C-X-C motif) ligand 9 | -1.594641026 | -3.020193587 | 10.49950515 | -10.3853962 | 1.35E-05 | 0.009806623 |
| Cmpk2 | TC1200000225.mm.2 | 22169 | cytidine monophosphate (UMP-CMP) kinase 2, mitochondrial | -2.094436297 | -4.270592661 | 8.056413678 | -10.38034499 | 1.36E-05 | 0.009806623 |
| Zbtb10 | TC0300000054.mm.2 | 229055 | zinc finger and BTB domain containing 10 | -1.551649251 | -2.931520714 | 8.565099318 | -10.3794199 | 1.36E-05 | 0.009806623 |

| | | | | | | | | | |
|---|---|---|---|---|---|---|---|---|---|
| Muc15 | TC0200001637.mm.2 | 269328 | mucin 15 | -1.275459549 | -2.420759152 | 7.159305488 | -10.31162694 | 1.42E-05 | 0.009806623 |
| Tgtp2 | TC1100002649.mm.2 | 100039796 | T cell specific GTPase 2 | -2.434781575 | -5.406824673 | 10.19495247 | -10.20527854 | 1.52E-05 | 0.009806623 |
| Ifit3b | TC1900000502.mm.2 | 667370 | interferon-induced protein with tetratricopeptide repeats 3B | -1.050899134 | -2.07182067 | 7.595583742 | -10.19086424 | 1.54E-05 | 0.009806623 |
| H2-Q2 | TC1700000682.mm.2 | 15013 | histocompatibility 2, Q region locus 2 | -1.471861414 | -2.773795473 | 9.342125702 | -10.1847925 | 1.54E-05 | 0.009806623 |
| Ifit1 | TC1900000504.mm.2 | 15957 | interferon-induced protein with tetratricopeptide repeats 1 | -1.606887549 | -3.045940054 | 8.395442755 | -10.04696838 | 1.69E-05 | 0.010214373 |
| Gm4951 | TC1800000606.mm.2 | 240327 | predicted gene 4951 | -2.479755439 | -5.578029016 | 9.681469039 | -10.03649854 | 1.70E-05 | 0.010214373 |
| Epsti1 | TC1400001086.mm.2 | 108670 | epithelial stromal interaction 1 (breast) | -1.004659483 | -2.006469857 | 8.981416546 | -9.70569229 | 2.14E-05 | 0.012123084 |
| Saa3 | TC0700003039.mm.2 | 20210 | serum amyloid A 3 | 1.615243909 | 3.063633904 | 10.05383323 | 9.607645145 | 2.29E-05 | 0.01237615 |
| F830016B08Rik | TC1800000609.mm.2 | 240328 | RIKEN cDNA F830016B08 gene | -1.88227536 | -3.686560313 | 8.768363779 | -9.600873931 | 2.30E-05 | 0.01237615 |
| S100a9 | TC0300002376.mm.2 | 20202 | S100 calcium binding protein A9 (calgranulin B) | 1.368853168 | 2.582651835 | 9.294831168 | 9.532732604 | 2.41E-05 | 0.012660442 |
| Psmb8 | TC1700000622.mm.2 | 16913 | proteasome (prosome, macropain) subunit, beta type 8 (large multifunctional peptidase 7) | -1.645720186 | -3.129040183 | 10.63112533 | -9.372193929 | 2.71E-05 | 0.013847909 |
| Prlr | TC1500002327.mm.2 | 19116 | prolactin receptor | -1.470082201 | -2.77037678 | 7.637792678 | -9.278952041 | 2.89E-05 | 0.013863596 |
| Phf11d | TC1400002869.mm.2 | 219132 | PHD finger protein 11D | -2.309983234 | -4.958773171 | 9.153501967 | -9.268738769 | 2.92E-05 | 0.013863596 |
| Sort1 | TC0300001137.mm.2 | 20661 | sortilin 1 | -1.425382751 | -2.685857476 | 9.549528405 | -9.195762017 | 3.07E-05 | 0.013863596 |
| Acot1 | TC1200000798.mm.2 | 26897 | acyl-CoA thioesterase 1 | -1.094426004 | -2.135281096 | 8.299875002 | -9.18368384 | 3.10E-05 | 0.013863596 |
| Klhl1 | TC1400002572.mm.2 | 93688 | kelch-like 1 | 1.273135664 | 2.416862946 | 7.993800615 | 9.181374301 | 3.11E-05 | 0.013863596 |

| | | | | | | | | | |
|---|---|---|---|---|---|---|---|---|---|
| Tmem139 | TC0600000455.mm.2 | 109218 | transmembrane protein 139 | -1.073571918 | -2.10463772 | 8.067020956 | -9.150377448 | 3.18E-05 | 0.013863596 |
| Hpn | TC0700002790.mm.2 | 15451 | hepsin | -1.118850785 | -2.171739083 | 8.60102842 | -9.1230121 | 3.24E-05 | 0.013863596 |
| Aldoc | TC1100001156.mm.2 | 11676 | aldolase C, fructose-bisphosphate | -1.64537949 | -3.128301339 | 9.582968004 | -9.108693349 | 3.28E-05 | 0.013863596 |
| Scrg1 | TC0800000627.mm.2 | 20284 | scrapie responsive gene 1 | -2.475786629 | -5.562705117 | 8.406915715 | -9.056095413 | 3.41E-05 | 0.013863596 |
| Trim30a | TC0700003943.mm.2 | 20128 | tripartite motif-containing 30A | -1.647638045 | -3.133202566 | 9.750626713 | -9.041976534 | 3.44E-05 | 0.013863596 |
| Alpl | TC0400003814.mm.2 | 11647 | alkaline phosphatase, liver/bone/kidney | -1.138543754 | -2.201586844 | 7.459138397 | -8.996250002 | 3.56E-05 | 0.013863596 |
| Tfap2c | TC0200002672.mm.2 | 21420 | transcription factor AP-2, gamma | -1.350890983 | -2.550696032 | 8.452283568 | -8.96889171 | 3.63E-05 | 0.013863596 |
| BC006965 | TC1100004023.mm.2 | 217294 | cDNA sequence BC006965 | -4.25079979 | -19.03786498 | 9.115368276 | -8.892964855 | 3.85E-05 | 0.014035341 |
| Fxyd3 | TC0700002789.mm.2 | 17178 | FXYD domain-containing ion transport regulator 3 | -1.135422511 | -2.196828902 | 11.17442269 | -8.88288996 | 3.88E-05 | 0.014035341 |
| Prl2c3 | TC1300001452.mm.2 | 18812 | prolactin family 2, subfamily c, member 3 | 1.104825067 | 2.15072798 | 7.612533492 | 8.873516146 | 3.90E-05 | 0.014035341 |
| Gm12185 | TC1100002643.mm.2 | 620913 | predicted gene 12185 | -2.463514356 | -5.515586702 | 10.343907 | -8.825031819 | 4.05E-05 | 0.014035341 |
| Gbp9 | TC0500003730.mm.2 | 236573 | guanylate-binding protein 9 | -1.33609488 | -2.524670095 | 8.925840839 | -8.821927701 | 4.06E-05 | 0.014035341 |
| H2-Q7 | TC1700002789.mm.2 | 15018 | histocompatibility 2, Q region locus 7 | -1.950527827 | -3.865159171 | 9.942267165 | -8.799721801 | 4.13E-05 | 0.014035341 |
| Clec2f | TC0600001525.mm.2 | 435921 | C-type lectin domain family 2, member f | -1.704054475 | -3.258153275 | 8.875033586 | -8.770956631 | 4.22E-05 | 0.014035341 |
| Cdkl5 | TC0X00003278.mm.2 | 382253 | cyclin-dependent kinase-like 5 | -1.311395137 | -2.481814242 | 9.69426716 | -8.735801018 | 4.33E-05 | 0.014035341 |

| | | | | | | | | | |
|---|---|---|---|---|---|---|---|---|---|
| Hrct1 | TC0400000450.mm.2 | 100039781 | histidine rich carboxyl terminus 1 | -1.533480751 | -2.894834259 | 9.291749769 | -8.730131819 | 4.35E-05 | 0.014035341 |
| Ifit1bl1 | TC1900001422.mm.2 | 667373 | interferon induced protein with tetratricpeptide repeats 1B like 1 | -1.335997054 | -2.524498909 | 7.115908436 | -8.702248706 | 4.44E-05 | 0.014035341 |
| Klk8 | TC0700000775.mm.2 | 259277 | kallikrein related-peptidase 8 | 1.16608063 | 2.244012372 | 9.0085304 | 8.672379121 | 4.55E-05 | 0.014035341 |
| Rsph9 | TC1700002164.mm.2 | 75564 | radial spoke head 9 homolog (Chlamydomonas) | -1.083809994 | -2.11962639 | 7.268369541 | -8.644552066 | 4.64E-05 | 0.014035341 |
| Lcn2 | TC0200003303.mm.2 | 16819 | lipocalin 2 | -1.441160566 | -2.715392155 | 12.26735052 | -8.63919715 | 4.66E-05 | 0.014035341 |
| Zbp1 | TC0200005290.mm.2 | 58203 | Z-DNA binding protein 1 | -1.376028944 | -2.595529605 | 9.430484718 | -8.634180985 | 4.68E-05 | 0.014035341 |
| Rangrf | TC1100003024.mm.2 | 57785 | RAN guanine nucleotide release factor | -1.266919458 | -2.406471701 | 8.398030454 | -8.57320051 | 4.91E-05 | 0.014284232 |
| Casp12 | TC0900000033.mm.2 | 12364 | caspase 12 | -1.238067168 | -2.358823003 | 8.847107191 | -8.543325508 | 5.02E-05 | 0.014284232 |
| Stat1 | TC0100000375.mm.2 | 20846 | signal transducer and activator of transcription 1 | -1.71938404 | -3.292957837 | 10.16845874 | -8.503869047 | 5.18E-05 | 0.014485098 |
| Pigr | TC0100001165.mm.2 | 18703 | polymeric immunoglobulin receptor | -3.597659673 | -12.10607822 | 8.39440803 | -8.47133171 | 5.31E-05 | 0.014485098 |
| Fam13a | TC0600002381.mm.2 | 58909 | family with sequence similarity 13, member A | -1.148452788 | -2.216760314 | 6.012432707 | -8.470679465 | 5.32E-05 | 0.014485098 |
| H2-T22 | TC1700002822.mm.2 | 15039 | histocompatibility 2, T region locus 22 | -1.241944756 | -2.365171432 | 10.45443003 | -8.414409768 | 5.56E-05 | 0.014802839 |
| Cd3g | TC0900002188.mm.2 | 12502 | CD3 antigen, gamma polypeptide | -1.996909255 | -3.991439807 | 7.826291384 | -8.401295426 | 5.61E-05 | 0.014802839 |
| Ehf | TC0200004224.mm.2 | 13661 | ets homologous factor | -2.489031391 | -5.614009057 | 10.22111735 | -8.392396222 | 5.65E-05 | 0.014802839 |
| Anpep | TC0700003570.mm.2 | 16790 | alanyl (membrane) aminopeptidase | -1.006931807 | -2.009632647 | 10.12717801 | -8.37860521 | 5.71E-05 | 0.014802839 |

| | | | | | | | | | |
|---|---|---|---|---|---|---|---|---|---|
| Oas2 | TC0500003175.mm.2 | 246728 | 2'-5' oligoadenylate synthetase 2 | -2.04963497 | -4.14001205 8 | 8.18610365 1 | -8.33383101 2 | 5.92E-05 | 0.01514950 9 |
| H2-Q5 | TC1700000685.mm.2 | 15016 | histocompatibility 2, Q region locus 5 | -1.57586262 6 | -2.98113689 4 | 9.28703131 5 | -8.28949889 4 | 6.13E-05 | 0.01536848 2 |
| Irx3 | TC0800002690.mm.2 | 16373 | Iroquois related homeobox 3 | -1.62677987 5 | -3.08822929 8 | 10.4902879 3 | -8.28545364 7 | 6.15E-05 | 0.01536848 2 |
| Ncoa7 | TC1000001947.mm.2 | 211329 | nuclear receptor coactivator 7 | -1.26760165 7 | -2.40760990 4 | 8.44152540 1 | -8.26693601 8 | 6.24E-05 | 0.01541360 9 |
| Chpt1 | TC1000002645.mm.2 | 212862 | choline phosphotransferase 1 | -1.94367760 6 | -3.84685008 | 8.63135991 1 | -8.25213455 6 | 6.32E-05 | 0.01541556 4 |
| Arl4a | TC1200001658.mm.2 | 11861 | ADP-ribosylation factor-like 4A | -1.14796765 5 | -2.21601501 2 | 8.48864867 5 | -8.22379830 4 | 6.46E-05 | 0.01558797 8 |
| Tnfsf10 | TC0300000161.mm.2 | 22035 | tumor necrosis factor (ligand) superfamily, member 10 | -2.51149908 4 | -5.70212270 1 | 7.68319643 7 | -8.08066509 5 | 7.25E-05 | 0.01680146 2 |
| Ifi203 | TC0100003891.mm.2 | 15950 | interferon activated gene 203 | -1.19729272 9 | -2.29308960 2 | 10.5768220 9 | -8.05825176 5 | 7.39E-05 | 0.01680146 2 |
| Wfdc3 | TC0200005127.mm.2 | 71856 | WAP four-disulfide core domain 3 | -2.96657535 9 | -7.81678501 | 8.87455955 | -8.05690511 6 | 7.40E-05 | 0.01680146 2 |
| Tap1 | TC1700000621.mm.2 | 21354 | transporter 1, ATP-binding cassette, sub-family B (MDR/TAP) | -1.66004955 4 | -3.16027379 5 | 9.78585485 1 | -8.05227250 3 | 7.42E-05 | 0.01680146 2 |
| Timd2 | TC1100002611.mm.2 | 171284 | T cell immunoglobulin and mucin domain containing 2 | -1.73904116 4 | -3.33813236 9 | 7.80333132 2 | -8.04865272 5 | 7.45E-05 | 0.01680146 2 |
| Cxcl10 | TC0500002756.mm.2 | 15945 | chemokine (C-X-C motif) ligand 10 | -1.26267303 6 | -2.39939891 7 | 7.98987821 2 | -7.99437381 7 | 7.78E-05 | 0.01704055 |
| Cd14 | TC1800001197.mm.2 | 12475 | CD14 antigen | -1.24037464 9 | -2.36259878 | 9.98555014 8 | -7.94122668 4 | 8.13E-05 | 0.01704055 |
| Nr3c2 | TC0800000854.mm.2 | 110784 | nuclear receptor subfamily 3, group C, member 2 | -2.03298195 7 | -4.09249869 5 | 7.81009866 2 | -7.92154984 6 | 8.27E-05 | 0.01704055 |
| Gbp11 | TC0500002896.mm.2 | 634650 | guanylate binding protein 11 | -1.03371507 3 | -2.04728942 8 | 8.43993688 8 | -7.88323961 1 | 8.53E-05 | 0.01704055 |

| | | | | | | | | | |
|---|---|---|---|---|---|---|---|---|---|
| Clcf1 | TC1900000035.mm.2 | 56708 | cardiotrophin-like cytokine factor 1 | 1.090803075 | 2.129925657 | 8.475858181 | 7.865304118 | 8.66E-05 | 0.01704055 |
| Enpp3 | TC1000001895.mm.2 | 209558 | ectonucleotide pyrophosphatase/phosphodiesterase 3 | -1.873309976 | -3.663721844 | 10.44861343 | -7.861176279 | 8.69E-05 | 0.01704055 |
| Scd1 | TC1900001562.mm.2 | 20249 | stearoyl-Coenzyme A desaturase 1 | -1.530349312 | -2.888557697 | 10.66574932 | -7.849302755 | 8.78E-05 | 0.01704055 |
| Atp13a4 | TC1600001443.mm.2 | 224079 | ATPase type 13A4 | -1.317086017 | -2.491623383 | 5.982541097 | -7.839744317 | 8.85E-05 | 0.01704055 |
| Csprs | TC1_GL456221_random00000019.mm.2 | 114564 | component of Sp100-rs | -2.83596767 | -7.14021576 | 8.622370551 | -7.809191105 | 9.08E-05 | 0.01704055 |
| Hey1 | TC0300001616.mm.2 | 15213 | hairy/enhancer-of-split related with YRPW motif 1 | -1.790470802 | -3.459277625 | 10.67040739 | -7.796052881 | 9.18E-05 | 0.01704055 |
| Gm4841 | TC1800001396.mm.2 | 225594 | predicted gene 4841 | -1.755574512 | -3.376607554 | 8.641062564 | -7.785518144 | 9.26E-05 | 0.01704055 |
| Irgm2 | TC1100004266.mm.2 | 54396 | immunity-related GTPase family M member 2 | -1.63202151 | -3.099469935 | 9.027096948 | -7.785494199 | 9.26E-05 | 0.01704055 |
| Slc5a8 | TC1000001085.mm.2 | 216225 | solute carrier family 5 (iodide transporter), member 8 | -2.975194246 | -7.863623484 | 8.925203576 | -7.776781639 | 9.33E-05 | 0.01704055 |
| Mapk6 | TC0900002676.mm.2 | 50772 | mitogen-activated protein kinase 6 | 1.070471817 | 2.100120075 | 8.830398679 | 7.775478422 | 9.34E-05 | 0.01704055 |
| Ifit2 | TC1900000500.mm.2 | 15958 | interferon-induced protein with tetratricopeptide repeats 2 | -1.31239019 | -2.483526585 | 8.51298408 | -7.771152584 | 9.37E-05 | 0.01704055 |
| Gbp7 | TC0300001445.mm.2 | 229900 | guanylate binding protein 7 | -1.28276894 | -2.433055012 | 10.33836581 | -7.76087396 | 9.45E-05 | 0.01704055 |
| Klhl13 | TC0X00001955.mm.2 | 67455 | kelch-like 13 | -1.027408657 | -2.038359696 | 7.945705301 | -7.756301618 | 9.49E-05 | 0.01704055 |
| Muc1 | TC0300000753.mm.2 | 17829 | mucin 1, transmembrane | -1.839916715 | -3.579893615 | 10.44818528 | -7.753278841 | 9.51E-05 | 0.01704055 |
| Coro2a | TC0400002603.mm.2 | 107684 | coronin, actin binding protein 2A | -1.644510949 | -3.126418585 | 9.442043835 | -7.74972328 | 9.54E-05 | 0.01704055 |
| Ephx2 | TC1400002266.mm.2 | 13850 | epoxide hydrolase 2, cytoplasmic | -1.197 | -2.292 | 7.277335443 | -7.744 | 9.58E-05 | 0.01704055 |

| | | | | | | | | | |
|---|---|---|---|---|---|---|---|---|---|
| | | | | 01081 | 64155 | | 393993 | | |
| Ntrk2 | TC1300000718.mm.2 | 18212 | neurotrophic tyrosine kinase, receptor, type 2 | -1.984226963 | -3.956506051 | 9.218999324 | -7.68600624 | 0.000100669 | 0.017712251 |
| Ttpa | TC0400000168.mm.2 | 50500 | tocopherol (alpha) transfer protein | -2.599613336 | -6.061241545 | 6.845919721 | -7.672804509 | 0.000101802 | 0.017712251 |
| Ptprf | TC0400003386.mm.2 | 19268 | protein tyrosine phosphatase, receptor type, F | -1.417277218 | -2.670809761 | 9.847436237 | -7.656305212 | 0.000103239 | 0.017712251 |
| Lbp | TC0200002426.mm.2 | 16803 | lipopolysaccharide binding protein | -2.13320791 | -4.386918517 | 9.805269123 | -7.649675815 | 0.000103822 | 0.017712251 |
| Igfbp2 | TC0100000610.mm.2 | 16008 | insulin-like growth factor binding protein 2 | -1.916075016 | -3.773949234 | 9.877017355 | -7.611909128 | 0.000107218 | 0.017988001 |
| Id2 | TC1200001544.mm.2 | 15902 | inhibitor of DNA binding 2 | -1.070801765 | -2.100600433 | 10.51165559 | -7.598229569 | 0.000108479 | 0.017988001 |
| Slc1a1 | TC1900000426.mm.2 | 20510 | solute carrier family 1 (neuronal/epithelial high affinity glutamate transporter, system Xag), member 1 | -1.128237627 | -2.185915491 | 7.579399572 | -7.589741692 | 0.00010927 | 0.017988001 |
| Tc2n | TC1200002241.mm.2 | 74413 | tandem C2 domains, nuclear | -1.512311307 | -2.852666921 | 8.137431577 | -7.584885816 | 0.000109725 | 0.017988001 |
| Irf7 | TC0700004530.mm.2 | 54123 | interferon regulatory factor 7 | -2.315408255 | -4.977454936 | 9.188636881 | -7.471418323 | 0.000120993 | 0.019381037 |
| D630039A03Rik | TC0400002729.mm.2 | 242484 | RIKEN cDNA D630039A03 gene | -1.08028616 | -2.114455444 | 8.557402234 | -7.436289983 | 0.000124741 | 0.019757564 |
| Oas1g | TC0500003180.mm.2 | 23960 | 2'-5' oligoadenylate synthetase 1G | -1.883159463 | -3.688820178 | 10.44036557 | -7.431822051 | 0.000125227 | 0.019757564 |
| Gbp2 | TC0300001447.mm.2 | 14469 | guanylate binding protein 2 | -1.341659382 | -2.5344266 | 8.725699088 | -7.389931836 | 0.000129888 | 0.020020833 |
| Slfn8 | TC1100003382.mm.2 | 276950 | schlafen 8 | -1.957163606 | -3.882978197 | 8.525230336 | -7.385277007 | 0.000130418 | 0.020020833 |
| Pik3r1 | TC1300002553.mm.2 | 18708 | phosphoinositide-3-kinase regulatory subunit 1 | -1.348014161 | -2.545614857 | 10.40096068 | -7.372624556 | 0.00013187 | 0.020020833 |
| Ifi44 | TC0300003133.mm.2 | 99899 | interferon-induced protein 44 | -2.347 | -5.090 | 9.847782641 | -7.366 | 0.00013262 | 0.020020833 |

| | | | | | | | | | |
|---|---|---|---|---|---|---|---|---|---|
| | | | | 765484 | 352216 | | 153811 | | |
| Phf11b | TC1400002162.mm.2 | 236451 | PHD finger protein 11B | -1.151448364 | -2.221367922 | 8.849388528 | -7.353782344 | 0.000134067 | 0.020094716 |
| Ddx60 | TC0800000659.mm.2 | 234311 | DEAD (Asp-Glu-Ala-Asp) box polypeptide 60 | -1.288650131 | -2.44299368 | 9.184276763 | -7.343909332 | 0.000135234 | 0.020125953 |
| Apol6 | TC1500000617.mm.2 | 71939 | apolipoprotein L 6 | -1.472887825 | -2.775769603 | 9.33521375 | -7.324698992 | 0.000137539 | 0.020192001 |
| Herc6 | TC0600003503.mm.2 | 67138 | hect domain and RLD 6 | -1.609100846 | -3.05061654 | 9.972979891 | -7.324172537 | 0.000137603 | 0.020192001 |
| H2-L | TC1700002790.mm.2 | 14980 | histocompatibility 2, D region locus L | -1.263801244 | -2.401276015 | 11.22189437 | -7.315125815 | 0.000138704 | 0.020212233 |
| Ubd | TC1700000761.mm.2 | 24108 | ubiquitin D | -1.105408247 | -2.151597544 | 6.058474708 | -7.297005347 | 0.000140939 | 0.020237328 |
| Ltf | TC0900001481.mm.2 | 17002 | lactotransferrin | -3.161666087 | -8.948625402 | 9.043082102 | -7.293430078 | 0.000141385 | 0.020237328 |
| Aqp5 | TC1500001018.mm.2 | 11830 | aquaporin 5 | -1.090339049 | -2.129240701 | 10.43197725 | -7.289722049 | 0.000141849 | 0.020237328 |
| Bcl2l15 | TC0300001062.mm.2 | 229672 | BCLl2-like 15 | -2.671451542 | -6.370698411 | 7.524126428 | -7.28024596 | 0.000143043 | 0.020237328 |
| Bmpr1b | TC0300003041.mm.2 | 12167 | bone morphogenetic protein receptor, type 1B | 1.005931593 | 2.008239861 | 6.167223051 | 7.275080308 | 0.000143698 | 0.020237328 |
| Trp53bp2 | TC0100001745.mm.2 | 209456 | transformation related protein 53 binding protein 2 | -1.063389425 | -2.089835556 | 10.11085629 | -7.24407987 | 0.000147703 | 0.020311639 |
| Isg15 | TC0400004171.mm.2 | 100038882 | ISG15 ubiquitin-like modifier | -1.503050063 | -2.834413149 | 8.277208579 | -7.215367353 | 0.000151525 | 0.020311639 |
| Rtp4 | TC1600000346.mm.2 | 67775 | receptor transporter protein 4 | -1.929174893 | -3.808373283 | 9.483566205 | -7.183767331 | 0.000155859 | 0.020699663 |
| Bst2 | TC0800002431.mm.2 | 69550 | bone marrow stromal cell antigen 2 | -1.134128246 | -2.194858974 | 11.90768423 | -7.175080218 | 0.000157075 | 0.020729939 |
| Clmn | TC1200002296.mm.2 | 94040 | calmin | -2.814320421 | -7.033878529 | 9.147619999 | -7.166510405 | 0.000158285 | 0.020759051 |

| | | | | | | | | | |
|---|---|---|---|---|---|---|---|---|---|
| Gabrp | TC1100002492.mm.2 | 216643 | gamma-aminobutyric acid (GABA) A receptor, pi | -2.25807338 | -4.78352248 7 | 7.804694087 | -7.151368133 | 0.000160448 | 0.020799365 |
| Il7 | TC0300001610.mm.2 | 16196 | interleukin 7 | -1.340889574 | -2.533074615 | 7.058414092 | -7.116762885 | 0.000165518 | 0.021178224 |
| Gbp4 | TC0500003731.mm.2 | 17472 | guanylate binding protein 4 | -1.545192128 | -2.918429318 | 9.75984505 | -7.090106471 | 0.000169546 | 0.021347643 |
| Greb1l | TC1800000076.mm.2 | 381157 | growth regulation by estrogen in breast cancer-like | -1.057558182 | -2.081405678 | 7.277201932 | -7.089331402 | 0.000169665 | 0.021347643 |
| Mndal | TC0100003890.mm.2 | 100040462 | myeloid nuclear differentiation antigen like | -1.154141494 | -2.225518501 | 11.37759893 | -7.087835305 | 0.000169894 | 0.021347643 |
| Boc | TC1600001643.mm.2 | 117606 | biregional cell adhesion molecule-related/down-regulated by oncogenes (Cdon) binding protein | -1.57345575 | -2.976167553 | 10.12208092 | -7.057312408 | 0.000174653 | 0.021685867 |
| Prl2c2 | TC1300001453.mm.2 | 18811 | prolactin family 2, subfamily c, member 2 | 1.18577364 | 2.274853501 | 7.460514934 | 7.040923243 | 0.000177269 | 0.021871271 |
| 1110008P14Rik | TC0200003302.mm.2 | 73737 | RIKEN cDNA 1110008P14 gene | -1.044785483 | -2.063059576 | 9.368757049 | -7.028557859 | 0.000179273 | 0.021871271 |
| AW112010 | TC1900001123.mm.2 | 107350 | expressed sequence AW112010 | -1.045739 | -2.064423559 | 10.01805763 | -7.020700641 | 0.000180559 | 0.021900852 |
| H2-K1 | TC1700001892.mm.2 | 14972 | histocompatibility 2, K1, K region | -1.257473062 | -2.390766216 | 12.2473726 | -7.010884718 | 0.00018218 | 0.021905361 |
| Gbp2b | TC0300001444.mm.2 | 14468 | guanylate binding protein 2b | -1.180366141 | -2.266342871 | 7.916522321 | -7.007855511 | 0.000182684 | 0.021905361 |
| Mgst1 | TC0600001639.mm.2 | 56615 | microsomal glutathione S-transferase 1 | -1.057155763 | -2.080825181 | 10.96077257 | -6.997699191 | 0.000184384 | 0.021943147 |
| Gbp3 | TC0300001446.mm.2 | 55932 | guanylate binding protein 3 | -1.662793173 | -3.166289507 | 10.23335546 | -6.992515386 | 0.000185258 | 0.021943147 |
| H2-Q4 | TC1700000684.mm.2 | 15015 | histocompatibility 2, Q region locus 4 | -1.209179352 | -2.312060824 | 8.006039542 | -6.98130074 | 0.000187166 | 0.021943147 |
| Morc4 | TC0X00003066.mm.2 | 75746 | microrchidia 4 | 1.026211449 | 2.036668882 | 9.117517788 | 6.966493473 | 0.000189719 | 0.021994858 |
| Slc4a11 | TC0200004636.mm.2 | 269356 | solute carrier family 4, sodium bicarbonate transporter-like, member 11 | 1.197720504 | 2.293769629 | 8.817401398 | 6.924681073 | 0.000197141 | 0.022300018 |

| | | | | | | | | | |
|---|---|---|---|---|---|---|---|---|---|
| Mmp15 | TC0800001133.mm.2 | 17388 | matrix metallopeptidase 15 | -1.687587807 | -3.22117671 | 9.175184006 | -6.92179844 | 0.000197665 | 0.022300018 |
| Cbr2 | TC1100004229.mm.2 | 12409 | carbonyl reductase 2 | -2.127018555 | -4.368138364 | 10.79940806 | -6.912769883 | 0.000199315 | 0.022365916 |
| Trim12c | TC0700003940.mm.2 | 319236 | tripartite motif-containing 12C | -1.035467413 | -2.049777637 | 7.822936689 | -6.885072614 | 0.000204474 | 0.022464281 |
| Casp1 | TC0900000031.mm.2 | 12362 | caspase 1 | -1.601287697 | -3.034140088 | 9.86055783 | -6.854362093 | 0.000210369 | 0.022991626 |
| St6galnac2 | TC1100004124.mm.2 | 20446 | ST6 (alpha-N-acetyl-neuraminyl-2,3-beta-galactosyl-1,3)-N-acetylgalactosaminide alpha-2,6-sialyltransferase 2 | -2.429802551 | -5.388196823 | 9.387460913 | -6.836289812 | 0.000213928 | 0.023020821 |
| Nme5 | TC1800001150.mm.2 | 75533 | NME/NM23 family member 5 | -1.387598528 | -2.616427945 | 8.813096247 | -6.824555287 | 0.000216274 | 0.023154592 |
| Gm8909 | TC1700002004.mm.2 | 667977 | predicted gene 8909 | -1.127840067 | -2.185313207 | 8.462705429 | -6.812641978 | 0.000218686 | 0.023176313 |
| Cfap47 | TC0X00002528.mm.2 | 636104 | cilia and flagella associated protein 47 | -1.085268495 | -2.121770322 | 5.528184494 | -6.801551604 | 0.000220958 | 0.023210057 |
| H2-Q8 | TC1700002788.mm.2 | 15019 | histocompatibility 2, Q region locus 8 | -1.485352965 | -2.799856651 | 8.884525913 | -6.800299382 | 0.000221217 | 0.023210057 |
| Irgm1 | TC1100002641.mm.2 | 15944 | immunity-related GTPase family M member 1 | -1.766753643 | -3.402873787 | 8.94672363 | -6.76654913 | 0.000228306 | 0.02371665 |
| Nlrc5 | TC0800001110.mm.2 | 434341 | NLR family, CARD domain containing 5 | -1.832576018 | -3.561724721 | 9.204955643 | -6.743576129 | 0.000233276 | 0.023910103 |
| Pde9a | TC1700000535.mm.2 | 18585 | phosphodiesterase 9A | -1.873727864 | -3.664783224 | 8.704008758 | -6.742161936 | 0.000233586 | 0.023910103 |
| Itga8 | TC0200002972.mm.2 | 241226 | integrin alpha 8 | -1.146273744 | -2.213414651 | 9.745332639 | -6.711664146 | 0.000240383 | 0.024101281 |
| Nmi | TC0200003528.mm.2 | 64685 | N-myc (and STAT) interactor | -1.188562616 | -2.279255436 | 9.651502403 | -6.707074592 | 0.000241425 | 0.024101281 |
| Elf5 | TC0200001532.mm.2 | 13711 | E74-like factor 5 | -1.824279695 | -3.541301542 | 7.969560086 | -6.703041016 | 0.000242345 | 0.024101281 |

| | | | | | | | | | |
|---|---|---|---|---|---|---|---|---|---|
| Ddx58 | TC0400002434.mm.2 | 230073 | DEAD (Asp-Glu-Ala-Asp) box polypeptide 58 | -1.995380704 | -3.987213073 | 9.382783454 | -6.688304213 | 0.0024574 | 0.024323575 |
| Cldn8 | TC1600002005.mm.2 | 54420 | claudin 8 | -2.464090493 | -5.517789778 | 8.112151965 | -6.678737387 | 0.00247972 | 0.024354026 |
| Ube2l6 | TC0200001222.mm.2 | 56791 | ubiquitin-conjugating enzyme E2L 6 | -1.275307414 | -2.420503892 | 8.804097946 | -6.663459737 | 0.00251583 | 0.024440843 |
| Sntb1 | TC1500001544.mm.2 | 20649 | syntrophin, basic 1 | -1.291399493 | -2.447653761 | 9.139872955 | -6.615810189 | 0.00263231 | 0.025261628 |
| Ephx1 | TC0100003679.mm.2 | 13849 | epoxide hydrolase 1, microsomal | -1.237795249 | -2.358378454 | 9.655232843 | -6.578725238 | 0.00272715 | 0.025777722 |
| Ccbe1 | TC1800001479.mm.2 | 320924 | collagen and calcium binding EGF domains 1 | 1.168556478 | 2.247866689 | 9.009745435 | 6.567862219 | 0.00275565 | 0.025898388 |
| Enpp5 | TC1700000842.mm.2 | 83965 | ectonucleotide pyrophosphatase/phosphodiesterase 5 | -2.06272828 | -4.17775613 | 8.662463228 | -6.564476667 | 0.0027646 | 0.025898388 |
| Trim34b | TC0700001520.mm.2 | 434218 | tripartite motif-containing 34B | -1.043006806 | -2.060517627 | 8.950106555 | -6.550296174 | 0.00280244 | 0.026136217 |
| Stat2 | TC1000001596.mm.2 | 20847 | signal transducer and activator of transcription 2 | -1.20716362 | -2.308832672 | 10.05655194 | -6.545595642 | 0.00281512 | 0.026138227 |
| Gstk1 | TC0600000454.mm.2 | 76263 | glutathione S-transferase kappa 1 | -1.191596972 | -2.284054336 | 8.552050782 | -6.535128638 | 0.00284356 | 0.026217949 |
| Prss22 | TC1700001612.mm.2 | 70835 | protease, serine 22 | 1.264389383 | 2.402255136 | 9.09406873 | 6.532007046 | 0.00285211 | 0.026217949 |
| Znfx1 | TC0200005181.mm.2 | 98999 | zinc finger, NFX1-type containing 1 | -1.30764492 | -2.475371261 | 8.498413424 | -6.515183347 | 0.00289867 | 0.026217949 |
| Htr2b | TC0100002736.mm.2 | 15559 | 5-hydroxytryptamine (serotonin) receptor 2B | 1.117748036 | 2.17007971 | 6.411299671 | 6.469084256 | 0.00303066 | 0.026720712 |
| Enpp4 | TC1700002135.mm.2 | 224794 | ectonucleotide pyrophosphatase/phosphodiesterase 4 | -1.808149012 | -3.501927005 | 9.135448479 | -6.449025308 | 0.00309018 | 0.027131488 |
| Calhm6 | TC1000001994.mm.2 | 215900 | calcium homeostasis modulator family member 6 | -1.369653174 | -2.584084369 | 8.376803825 | -6.378405909 | 0.00331036 | 0.028123317 |
| Dcxr | TC1100004228.mm.2 | 67880 | dicarbonyl L-xylulose reductase | -1.136068734 | -2.197813142 | 10.6327542 | -6.356072692 | 0.0033836 | 0.028480427 |

| | | | | | | | | | |
|---|---|---|---|---|---|---|---|---|---|
| Mmp19 | TC1000001621.mm.2 | 58223 | matrix metallopeptidase 19 | 1.242901606 | 2.366740623 | 8.284635399 | 6.349145687 | 0.000340668 | 0.028480427 |
| Tmem100 | TC1100001395.mm.2 | 67888 | transmembrane protein 100 | 1.342819113 | 2.536464753 | 7.606997741 | 6.328734437 | 0.000347573 | 0.02887435 |
| Vtcn1 | TC0300001006.mm.2 | 242122 | V-set domain containing T cell activation inhibitor 1 | -1.812738419 | -3.513084843 | 9.030947861 | -6.327101444 | 0.000348132 | 0.02887435 |
| Gm7609 | TC0100000743.mm.2 | 665378 | predicted pseudogene 7609 | -1.901196021 | -3.735227255 | 8.642350411 | -6.314534736 | 0.000352469 | 0.029118938 |
| Ifih1 | TC0200003630.mm.2 | 71586 | interferon induced with helicase C domain 1 | -1.002889152 | -2.004009228 | 8.836794591 | -6.291428491 | 0.000360601 | 0.029136733 |
| Cd3d | TC0900000529.mm.2 | 12500 | CD3 antigen, delta polypeptide | -1.534303329 | -2.896485271 | 7.858484232 | -6.290128169 | 0.000361065 | 0.029136733 |
| Usp18 | TC0600001369.mm.2 | 24110 | ubiquitin specific peptidase 18 | -1.238174489 | -2.35899848 | 9.82870286 | -6.284224903 | 0.000363179 | 0.029136733 |
| Sp140 | TC0100000765.mm.2 | 434484 | Sp140 nuclear body protein | -1.414701385 | -2.666045469 | 9.055311878 | -6.283119506 | 0.000363577 | 0.029136733 |
| Fezf1 | TC0600001930.mm.2 | 73191 | Fez family zinc finger 1 | -1.1058794 | -2.152300324 | 5.813397627 | -6.277642974 | 0.000365553 | 0.029136733 |
| Rsad2 | TC1200001552.mm.2 | 58185 | radical S-adenosyl methionine domain containing 2 | -1.539952329 | -2.907848949 | 7.45340503 | -6.272105582 | 0.000367563 | 0.029136733 |
| Akr1c18 | TC1300001377.mm.2 | 105349 | aldo-keto reductase family 1, member C18 | 1.192838994 | 2.286021534 | 7.36524762 | 6.271778048 | 0.000367682 | 0.029136733 |
| Pcsk5 | TC1900001219.mm.2 | 18552 | proprotein convertase subtilisin/kexin type 5 | 1.271090803 | 2.413439737 | 9.582440254 | 6.271462831 | 0.000367797 | 0.029136733 |
| Nqo1 | TC0800002877.mm.2 | 18104 | NAD(P)H dehydrogenase, quinone 1 | -1.010590976 | -2.014736234 | 7.561370695 | -6.271315844 | 0.000367851 | 0.029136733 |
| Plau | TC1400000176.mm.2 | 18792 | plasminogen activator, urokinase | 1.185333388 | 2.274159414 | 9.715561585 | 6.262390503 | 0.00037112 | 0.029211331 |
| Arhgef38 | TC0300002977.mm.2 | 77669 | Rho guanine nucleotide exchange factor (GEF) 38 | -1.42977368 | -2.694044497 | 8.169347413 | -6.2608575 | 0.000371684 | 0.029211331 |
| Kit | TC0500000710.mm.2 | 16590 | KIT proto-oncogene receptor tyrosine kinase | -2.21504511 | -4.642960831 | 10.36979008 | -6.233010793 | 0.000382112 | 0.029918794 |

| | | | | | | | | | |
|---|---|---|---|---|---|---|---|---|---|
| Gzmb | TC1400002120.mm.2 | 14939 | granzyme B | -1.048004454 | -2.067667856 | 7.889902389 | -6.222115652 | 0.00038628 | 0.030132739 |
| Rgs20 | TC0100001914.mm.2 | 58175 | regulator of G-protein signaling 20 | -1.355567695 | -2.558977912 | 6.876674835 | -6.201621789 | 0.000394259 | 0.030509447 |
| Prom1 | TC0500002320.mm.2 | 19126 | prominin 1 | -1.864625802 | -3.641734638 | 7.247931542 | -6.19487798 | 0.000396925 | 0.030509447 |
| Palmd | TC0300002815.mm.2 | 114301 | palmdelphin | -1.042886389 | -2.060345651 | 10.25009432 | -6.169725201 | 0.000407047 | 0.031059895 |
| Stac2 | TC1100003665.mm.2 | 217154 | SH3 and cysteine rich domain 2 | -1.227721062 | -2.341967511 | 8.123847454 | -6.148413221 | 0.000415849 | 0.031276594 |
| Epb41l4b | TC0400002713.mm.2 | 54357 | erythrocyte membrane protein band 4.1 like 4b | -1.342677556 | -2.536215889 | 9.674318739 | -6.136300635 | 0.000420946 | 0.031546881 |
| Mx2 | TC1600001092.mm.2 | 17858 | MX dynamin-like GTPase 2 | -1.408428521 | -2.65447862 | 8.449560916 | -6.121934334 | 0.000427082 | 0.031892832 |
| Sox10 | TC1500001863.mm.2 | 20665 | SRY (sex determining region Y)-box 10 | -2.322848703 | -5.003191603 | 8.810457852 | -6.106677722 | 0.000433708 | 0.032158736 |
| Optn | TC0200002872.mm.2 | 71648 | optineurin | -1.312914237 | -2.48442887 | 7.126136497 | -6.095456223 | 0.000438654 | 0.032410995 |
| Tspan12 | TC0600001917.mm.2 | 269831 | tetraspanin 12 | -1.304831912 | -2.470549414 | 8.904980063 | -6.075383673 | 0.00044766 | 0.032730687 |
| Cdc42ep3 | TC1700002564.mm.2 | 260409 | CDC42 effector protein (Rho GTPase binding) 3 | -1.003484332 | -2.004836148 | 10.35488017 | -6.063257564 | 0.000453201 | 0.032906475 |
| Ifit1bl2 | TC1900001423.mm.2 | 112419 | interferon induced protein with tetratricopeptide repeats 1B like 2 | -1.243830265 | -2.368264577 | 6.306451654 | -6.048574782 | 0.000460012 | 0.033084467 |
| Gzma | TC1300002691.mm.2 | 14938 | granzyme A | -1.183652424 | -2.271511208 | 6.59335336 | -6.04778436 | 0.000460382 | 0.033084467 |
| Rap1gap | TC0400001712.mm.2 | 110351 | Rap1 GTPase-activating protein | -1.110759843 | -2.159593595 | 8.9653893 | -5.999658497 | 0.000483542 | 0.034153711 |
| Hmga1 | TC1700000461.mm.2 | 15361 | high mobility group AT-hook 1 | 1.200597427 | 2.2983487 | 10.1124709 | 5.964628001 | 0.000501218 | 0.03471138 |

| | | | | | | | | | |
|---|---|---|---|---|---|---|---|---|---|
| Sp110 | TC1_GL456212_random00000013.mm.2 | 109032 | Sp110 nuclear body protein | -1.475112856 | -2.780053903 | 7.993216478 | -5.957604224 | 0.000504847 | 0.034717968 |
| Chrnb1 | TC1100003063.mm.2 | 11443 | cholinergic receptor, nicotinic, beta polypeptide 1 (muscle) | 1.077819313 | 2.110843051 | 8.752253915 | 5.956409803 | 0.000505468 | 0.034717968 |
| Trim14 | TC0400002601.mm.2 | 74735 | tripartite motif-containing 14 | -1.685651904 | -3.216857221 | 8.501667835 | -5.954855312 | 0.000506276 | 0.034717968 |
| Mfsd4a | TC0100003068.mm.2 | 213006 | major facilitator superfamily domain containing 4A | -1.014038306 | -2.019556218 | 8.685663541 | -5.936307182 | 0.000516034 | 0.035051443 |
| Itga3 | TC1100003584.mm.2 | 16400 | integrin alpha 3 | 1.125239961 | 2.18137826 | 10.09836075 | 5.932631298 | 0.000517993 | 0.035051443 |
| Ptpn22 | TC0300001064.mm.2 | 19260 | protein tyrosine phosphatase, non-receptor type 22 (lymphoid) | -1.573060308 | -2.975351899 | 8.175339216 | -5.929658899 | 0.000519583 | 0.035051443 |
| Sox6 | TC0700004148.mm.2 | 20679 | SRY (sex determining region Y)-box 6 | -2.281081013 | -4.860420093 | 7.334046876 | -5.926718223 | 0.000521161 | 0.035051443 |
| Cdhr1 | TC1400001745.mm.2 | 170677 | cadherin-related family member 1 | -1.496870257 | -2.822297863 | 7.735333132 | -5.914545667 | 0.000527752 | 0.035156635 |
| Mcc | TC1800001280.mm.2 | 328949 | mutated in colorectal cancers | -1.219584748 | -2.328796777 | 8.320496356 | -5.89283528 | 0.000539737 | 0.0358413 |
| Plekhs1 | TC1900000800.mm.2 | 226245 | pleckstrin homology domain containing, family S member 1 | -1.369817739 | -2.584379146 | 8.668039885 | -5.878852595 | 0.000547617 | 0.036249859 |
| Sp100 | TC0100000766.mm.2 | 20684 | nuclear antigen Sp100 | -1.417311269 | -2.670872799 | 9.823949958 | -5.844446023 | 0.000567559 | 0.037101734 |
| Trpm6 | TC1900000327.mm.2 | 225997 | transient receptor potential cation channel, subfamily M, member 6 | -2.918097059 | -7.558484818 | 8.437918676 | -5.835188527 | 0.000573062 | 0.037290879 |
| Mme | TC0300000500.mm.2 | 17380 | membrane metallo endopeptidase | -2.649944356 | -6.2764307 | 8.712741077 | -5.831347536 | 0.000575362 | 0.037290879 |
| Krt23 | TC1100003709.mm.2 | 94179 | keratin 23 | -1.66791167 | -3.177543047 | 8.010251822 | -5.815755206 | 0.000584807 | 0.037758749 |
| Car2 | TC0300000096.mm.2 | 12349 | carbonic anhydrase 2 | -1.188711021 | -2.279489907 | 8.140198582 | -5.796648525 | 0.000596617 | 0.038325436 |
| Tfec | TC0600001890.mm.2 | 21426 | transcription factor EC | 1.241746481 | 2.364846399 | 9.521485299 | 5.767964508 | 0.000614849 | 0.038861422 |

| | | | | | | | | | |
|---|---|---|---|---|---|---|---|---|---|
| Lurap1l | TC0400000811.mm.2 | 52829 | leucine rich adaptor protein 1-like | -1.931963523 | -3.815741724 | 8.477996711 | -5.763052548 | 0.000618033 | 0.038945352 |
| Lgi1 | TC1900000547.mm.2 | 56839 | leucine-rich repeat LGI family, member 1 | 1.030903615 | 2.04330365 | 5.950748286 | 5.754420627 | 0.000623673 | 0.039066104 |
| Gvin1 | TC0700004002.mm.2 | 74558 | GTPase, very large interferon inducible 1 | -1.452926633 | -2.737628402 | 9.279465832 | -5.731960717 | 0.000638617 | 0.039150651 |
| Plaur | TC0700000383.mm.2 | 18793 | plasminogen activator, urokinase receptor | 1.014466821 | 2.020156163 | 9.525169729 | 5.724577846 | 0.000643616 | 0.039150651 |
| Thrsp | TC0700003764.mm.2 | 21835 | thyroid hormone responsive | -2.235822942 | -4.710313039 | 8.092344774 | -5.716993873 | 0.000648797 | 0.039150651 |
| Ccdc141 | TC0200003809.mm.2 | 545428 | coiled-coil domain containing 141 | -1.23461534 | -2.35318597 | 8.882312677 | -5.707572956 | 0.000655297 | 0.039400447 |
| Lifr | TC1500000045.mm.2 | 16880 | LIF receptor alpha | -1.175345156 | -2.258469071 | 8.451213746 | -5.665974782 | 0.000684879 | 0.040597457 |
| Wfdc18 | TC1100001265.mm.2 | 14038 | WAP four-disulfide core domain 18 | -3.017221865 | -8.096070567 | 11.75445042 | -5.638911969 | 0.000704921 | 0.041530618 |
| Slpi | TC0200005112.mm.2 | 20568 | secretory leukocyte peptidase inhibitor | -1.223889831 | -2.335756416 | 11.62247771 | -5.632269247 | 0.000709939 | 0.04161275 |
| Atp6v1b1 | TC0600000937.mm.2 | 110935 | ATPase, H+ transporting, lysosomal V1 subunit B1 | -1.363240017 | -2.57262293 | 9.143102518 | -5.625403735 | 0.000715168 | 0.041690025 |
| Shf | TC0200004508.mm.2 | 435684 | Src homology 2 domain containing F | -1.006995997 | -2.009722065 | 8.867801744 | -5.605572292 | 0.000730513 | 0.042462842 |
| C1qtnf1 | TC1100001988.mm.2 | 56745 | C1q and tumor necrosis factor related protein 1 | -1.517601851 | -2.863147213 | 10.04989756 | -5.597246353 | 0.000737064 | 0.042725299 |
| 5330417C22Rik | TC0300002743.mm.2 | 229722 | RIKEN cDNA 5330417C22 gene | -2.543622432 | -5.830511411 | 7.321156554 | -5.581303517 | 0.000749792 | 0.043105857 |
| Pipox | TC1100003276.mm.2 | 19193 | pipecolic acid oxidase | -1.404018601 | -2.646377 | 7.354294416 | -5.574710454 | 0.000755127 | 0.043215678 |
| Dlx5 | TC0600001834.mm.2 | 13395 | distal-less homeobox 5 | -1.154543586 | -2.226138859 | 8.453203119 | -5.573856046 | 0.000755821 | 0.043215678 |
| Gm18852 | TC0700004003.mm.2 | 100417829 | GTPase, very large interferon inducible 1 pseudogene | -1.307 | -2.475 | 9.867740694 | -5.561 | 0.000765994 | 0.043604162 |

| | | | | | | | | | |
|---|---|---|---|---|---|---|---|---|---|
| | | | | 79411 | 627255 | | 438611 | | |
| Slc28a3 | TC1300002102.mm.2 | 114304 | solute carrier family 28 (sodium-coupled nucleoside transporter), member 3 | -2.194568094 | -4.577526042 | 8.730311683 | -5.557986012 | 0.00076885 | 0.043604162 |
| Pacsin1 | TC1700000464.mm.2 | 23969 | protein kinase C and casein kinase substrate in neurons 1 | -1.25290559 | -2.383209187 | 6.506440074 | -5.530148348 | 0.00079231 | 0.044573271 |
| S100a8 | TC0300000811.mm.2 | 20201 | S100 calcium binding protein A8 (calgranulin A) | 1.092641919 | 2.132642168 | 8.953163547 | 5.46532876 | 0.000850081 | 0.046332721 |
| Homer2 | TC0700003614.mm.2 | 26557 | homer scaffolding protein 2 | -1.340650331 | -2.532654589 | 7.873486883 | -5.457330343 | 0.000857528 | 0.046497085 |
| Gdpd1 | TC1100003466.mm.2 | 66569 | glycerophosphodiester phosphodiesterase domain containing 1 | -1.06477653 | -2.091845832 | 9.496487471 | -5.350683644 | 0.00096407 | 0.049950735 |
| Il1rl1 | TC0100000298.mm.2 | 17082 | interleukin 1 receptor-like 1 | 1.40808741 | 2.653851067 | 7.74132431 | 5.344882198 | 0.000970274 | 0.049997878 |
| Il33 | TC1900000448.mm.2 | 77125 | interleukin 33 | -1.062819944 | -2.08901079 | 9.500914507 | -5.339473825 | 0.000976097 | 0.049997878 |
| Bhlha15 | TC0500001749.mm.2 | 17341 | basic helix-loop-helix family, member a15 | -1.016090727 | -2.022431343 | 6.598313456 | -5.338976852 | 0.000976634 | 0.049997878 |
| Acsbg1 | TC0900002322.mm.2 | 94180 | acyl-CoA synthetase bubblegum family member 1 | 1.150316483 | 2.219625808 | 9.371989297 | 5.338736578 | 0.000976893 | 0.049997878 |
